# A tracheal aspirate-derived airway basal cell model reveals a proinflammatory epithelial defect in congenital diaphragmatic hernia

**DOI:** 10.1101/2022.11.10.515365

**Authors:** Richard Wagner, Gaurang M. Amonkar, Wei Wang, Jessica E. Shui, Kamakshi Bankoti, Wai Hei Tse, Frances A. High, Jill M. Zalieckas, Terry L. Buchmiller, Augusto Zani, Richard Keijzer, Patricia K. Donahoe, Paul H. Lerou, Xingbin Ai

**Affiliations:** Division of Newborn Medicine, Department of Pediatrics, Massachusetts General Hospital, Boston, MA, USA; Pediatric Surgical Research Laboratories, Department of Surgery, Massachusetts General Hospital, Harvard Medical School, Boston, MA, USA; Department of Pediatric Surgery, University Hospital Leipzig, Leipzig, Sachsen, Germany; Departments of Surgery, Pediatrics & Child Health, Physiology & Pathophysiology, University of Manitoba and Children’s Hospital Research Institute of Manitoba, Winnipeg, Manitoba, Canada; Boston Children’s Hospital, Harvard Medical School, Boston, MA, USA; Division of Medical Genetics, Department of Pediatrics, Massachusetts General Hospital, Boston, MA, USA; Division of Pediatric Surgery, Department of Surgery, Boston Children’s Hospital, Boston, MA, USA; Department of Pediatric Surgery, University of Toronto, Hospital for Sick Children, Toronto, Canada

## Abstract

**Rationale:** Congenital diaphragmatic hernia (CDH) is characterized by incomplete closure of the diaphragm and lung hypoplasia. The pathophysiology of lung defects in CDH is poorly understood.

**Objectives:** To establish a translational model of human airway epithelium in CDH for pathogenic investigation and therapeutic testing.

**Methods:** We developed a robust methodology of epithelial progenitor derivation from tracheal aspirates of newborns. Basal stem cells (BSCs) from CDH patients and preterm and term, non-CDH controls were derived and analyzed by bulk RNA-sequencing, ATAC-sequencing, and air-liquid-interface differentiation. Lung sections from fetal human CDH samples and the nitrofen rat model of CDH were subjected to histological assessment of epithelial defects. Therapeutics to restore epithelial differentiation were evaluated in human epithelial cell culture and the nitrofen rat model of CDH.

**Measurements and Main Results:** Transcriptomic and epigenetic profiling of CDH and non-CDH basal stem cells reveals a disease-specific, proinflammatory signature independent of severity or hernia size. In addition, CDH basal stem cells exhibit defective epithelial differentiation *in vitro* that recapitulates epithelial phenotypes found in fetal human CDH lung samples and fetal tracheas of the nitrofen rat model of CDH. Furthermore, steroid treatment normalizes epithelial differentiation phenotypes of human CDH basal stem cells *in vitro* and in nitrofen rat tracheas *in vivo*.

**Conclusions:** Our findings have identified an underlying proinflammatory signature and BSC differentiation defects as a potential therapeutic target for airway epithelial defects in CDH.

## INTRODUCTION

Congenital diaphragmatic hernia (CDH) affects 1 out of 3000 live births and is characterized by abnormal development of lung and diaphragm (1, 2). Lung hypoplasia in CDH features reduced airway branching, impaired alveolarization, and thickened mesenchyme (3, 4). Surgical repair of the diaphragm and current non-surgical treatments are ineffective to restore lung structure and function in CDH causing significant mortality and long-term complications in survivors (5–7).

The etiology of lung defects in CDH is complex. In addition to mechanical compression that affects lung branching and epithelial formation (8–11), pathogenic mechanisms inherent to lung development also contribute to lung hypoplasia in CDH (12). For example, a few CDH-associated genes play primary roles in the formation of airway epithelium and lung mesenchyme (13–18). In an established CDH model induced by oral nitrofen administration to pregnant rat dams, lung abnormalities in exposed fetuses are evident before the closure of the diaphragm (3, 19). Despite these findings in animal models, pathogenic changes in the human CDH lung remain poorly characterized due to limited access to neonatal lung tissue. Lung biopsy at the time of surgery is contraindicated and postmortem lung tissues are rare and often affected by artifacts from clinical procedures and tissue handling (20). To directly study lung defects in human CDH, translational models of patient-specific, epithelium and mesenchyme are needed.

BSCs are multipotent progenitor cells localized along the entire conducting airways in humans (21, 22) while restricted to the trachea and main bronchi in rodents (23). BSCs can self-renew and give rise to multiple epithelial cell types during development, homeostasis and regeneration following injury (24, 25). Importantly during development, P63^+^ lineage labeled epithelial progenitors can generate both proximal and distal epithelium (24). A useful *in vitro* model of functional airway epithelium is air-liquid-interface (ALI) culture in which BSCs differentiate into multiple epithelial cell types (26, 27).

Tracheal aspirates (TA) can be collected during routine care of intubated patients and are usually considered medical waste. TA contain epithelial and mesenchymal progenitors and provide a renewable surrogate of neonatal lung tissue (28–31).

To investigate epithelial defects in CDH patients, we derived epithelial progenitors from TA of CDH newborns and preterm and term, non-CDH controls. Using a combination of transcriptomic and epigenetic profiling, *in vitro* assays, histological characterization of fetal human CDH lung samples and the nitrofen rat CDH model, we have identified a proinflammatory signature of CDH BSCs that correlates with differentiation defects of the airway epithelium in CDH.

## MATERIALS AND METHODS

### Human samples

Patient enrollment, TA sample acquisition, and experiments were approved by the Institutional Review Board at Massachusetts General Hospital (IRB #2019P003296, PI: Lerou) and Boston Children’s Hospital (IRB #2000P000372, PI: High). Demographic patient data is summarized in Table 1 (CDH patients) and Supplementary Table 1 (non-CDH control patients). Human fetal lung sections were obtained from a previously established tissue bank at the University of Manitoba (IRB protocol #HS15293, PI: Keijzer) (20).

**Table 1:** Demographic and clinical information of CDH patients from whom TA BSCs were derived and the summary of assays performed for each CDH BSC line.

| Patient # | Sex | Gestational Age (weeks) | Survival (Y/N) | Side | Defect Size | ECMO (Y/N) | Comorbidities | Differentiated (Y/N) | Assay |
| --- | --- | --- | --- | --- | --- | --- | --- | --- | --- |
| 1 | M | 38 | Y | L | C type | N | Mild coarctation | Y | R, A, D |
| 2 | M | 39 | Y | L | C type | Y | Isolated | Y | R, A, D |
| 3 | M | 39 | Y | R | A type | N | VAWH, BAV | N | R, D |
| 4 | M | 28 | Y | R | B type | N | Bilateral grade II IVH with cystic changes | N | R, D |
| 5 | M | 39 | Y | L | B type | N | Isolated | Y | R, D |
| 6 | M | 37 | Y | L | C type | Y | Isolated | Y | R, D |
| 7 | M | 36 | Y | L | C type | Y | Hypospadia, cryptorchidism, camptodactyly, IUGR | N | R, D |
| 8 | F | 37 | Y | L | C type | N | Isolated | Y | R, D |
| 9 | M | 37 | Y | L | C type | Y | Isolated | N | R, D |
| 10 | M | 39 | Y | L | C type | Y | Isolated | Y | R, D |
| 11 | M | 39 | Y | L | D type | Y | Isolated | Y | R, D |
| 12 | M | 37 | Y | R | D type | Y | Isolated | N | R, D |
| 13 | M | 38 | Y | R | B type | Y | Tetralogy of Fallot | N | D |

\***VAWH**: ventral abdominal wall hernia, **BAV**: bicuspid aortic valve, **IUGR**: intrauterine growth restriction, **PFO**: patent foramen ovale, **MTR**: moderate tricuspid regurgitation
\***R** = RNA Sequencing, **A** = ATAC Sequencing, **D** = Differentiation

### Nitrofen rat model of CDH

Animal work was approved by the Institutional Animal Care and Use Committee at Massachusetts General Hospital (IACUC #2022N000003, PI: Ai). Sprague-Dawley rat dams (Charles River Laboratory) at E9.5 were orally gavaged with nitrofen (100mg in 1 mL olive oil) or olive oil (vehicle) (32). Rat fetuses were harvested at E21.5.

### Immunohistochemistry

Tracheas and lungs were fixed in 4% paraformaldehyde/PBS and processed for paraffin embedding and sectioning before immunohistochemistry using standard staining procedures.

### BSC derivation, expansion, and differentiation assays

A detailed protocol of TA BSC derivation, expansion, differentiation in ALI, and staining of ALI cultures for epithelial cell markers was recently published (28, 29). For DXM treatment (Cat#D4902, Sigma Aldrich), BSCs were treated for on alternate days during 7-day cell expansion and for 7-21 days during ALI at 10μM and 10mM. For Poly (I:C) treatment (10μM, Cat#tlrl-pic, Invivogen), BSCs were treated every 2 days during 7-day cell expansion followed by treatment at days 3, 5, 7, and 14 in ALI. For treatment with a specific NF-κB inhibitor JSH-23 (10μM, Cat#S7351, Selleck Chemicals) or STAT3 inhibitor S3I-201 (20μM, Cat#S1155, Selleck Chemicals), ALI cultures were treated daily during the first week of differentiation. Bulk RNA-seq and ATAC seq of BSCs were performed by ActiveMotif.inc (Carlsbad, CA, USA).

### Statistical Analysis and data availability

R-Studio and GraphPad Prism 6 were used for data analyses. Data in all graphs represent mean ± SEM from minimally 3 independent experiments. For comparisons between two or more conditions, statistical significance was analyzed using Mann-Whitney U test, Student’s t test or Kruskall-Wallis test when appropriate. Statistical significance was considered if p-value was less than 0.05. Only comparisons that reached statistical significance were highlighted by horizontal bars in graphs. For histological quantification, each dot represents quantification of an average of 4 sections per sample. All sequencing data can be accessed at NCBI Gene Accession Omnibus via accession number GSE211790.

More details of Materials & Methods are provided in Supplementary Materials.

## RESULTS

### BSCs derived from TA are *bona fide* lower respiratory epithelial progenitors

We collected TA from newborns within the first 24-48 hours of intubation for BSC derivation to minimize the impact of ventilation and postnatal clinical course on cellular phenotypes (Fig. 1A). To establish that TA epithelial progenitors are BSCs in the lower respiratory tract, we showed that TA contained rare P63^+^ BSCs (Fig. 1A). All TA BSCs proliferated to form clones after one week in culture (Fig. 1B) and reached confluency within 3-4 weeks as passage 0 (P0). BSCs were collected at P2 for assays and cryopreservation in liquid nitrogen. Our pipeline from TA collection to BSC derivation has achieved a success rate of almost 100% per TA sample. In this study, bulk RNA-seq and ATAC-seq were performed at P2 and all other assays were conducted at P3-P4. Antibody staining and transcriptomic assays showed that all BSCs derived from TA of preterm and term newborns expressed basal cell markers, such as P63 and KRT5, and NKX2.1, the lineage-specifying transcription factor of lower respiratory epithelial cells (21) (Fig. 1C). In contrast, BSCs derived from nasopharyngeal swabs (NP BSCs) lacked NKX2.1 (Fig. 1C) and exhibited significant differences in gene expression compared to TA BSCs (Fig S1). These findings, together with our previous publication that TA BSCs can differentiate into multiple airway epithelial cell types (29), indicate that TA BSCs are *bona fide* lower respiratory epithelial progenitors.

**Fig. 1.**
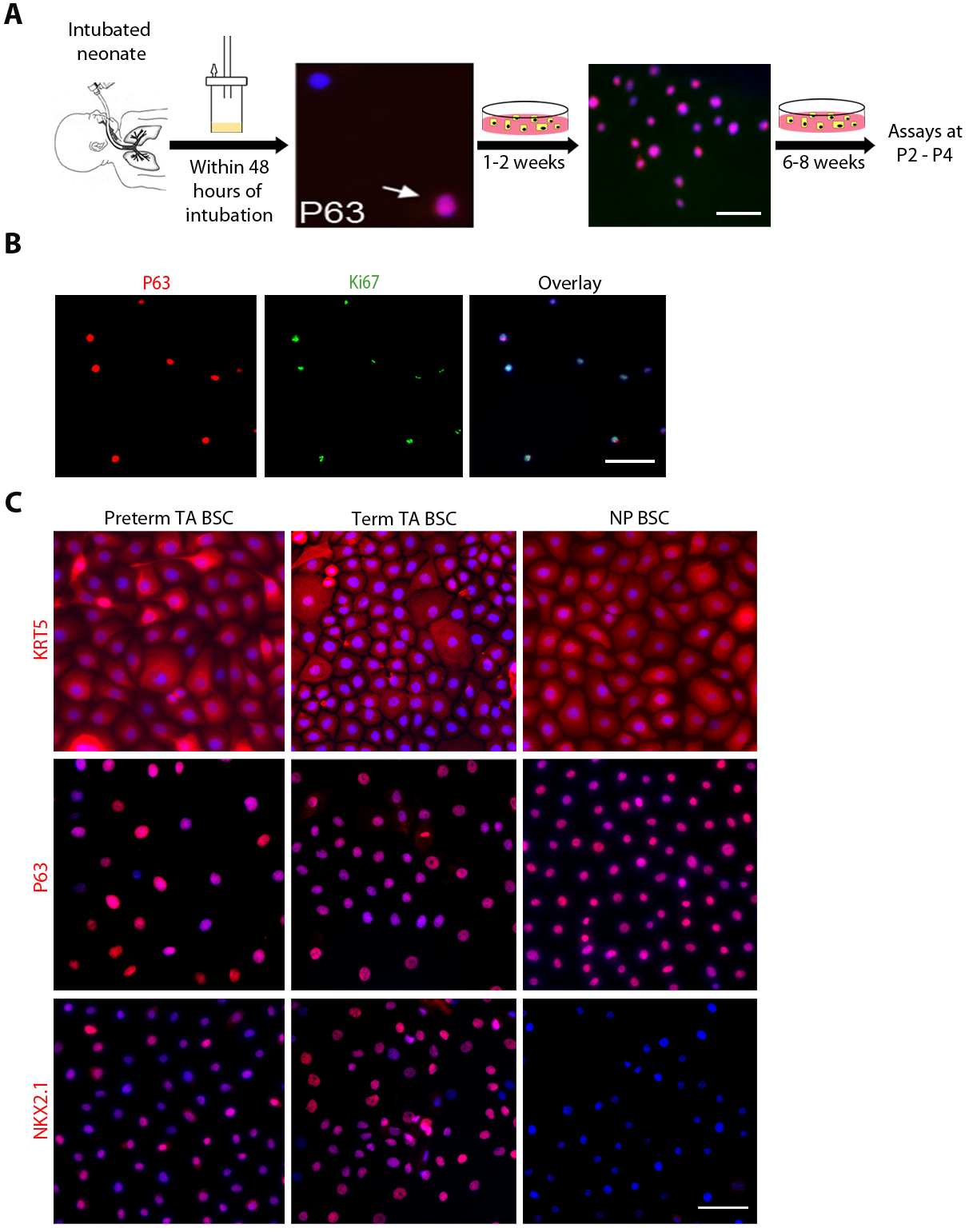
TA BSCs are *bona fide* lower respiratory epithelial progenitor cells. (**A**) Schematic of the workflow. The images show P63 staining of fresh neonatal TA sample (arrow) and the TA sample culture after 1-2 weeks. Cultured BSCs reached confluency within 3-4 weeks as passage 0 (P0). Bulk RNA-seq and ATAC-seq of BSCs were performed at P2 and all other assays were conducted at P3-P4. (**B**) Co-staining for P63 and Ki67 of BSC clones after 1-2 weeks in culture of TA samples. (**C**) Representative images of KRT5, P63, and NKX2.1 staining of confluent cultures of BSCs derived from TA of term and preterm newborns and nasopharyngeal (NP) samples. Scale bars, 25 μM.

### Transcriptome profiling identifies a proinflammatory signature of CDH BSCs

We derived TA BSCs from 13 CDH newborns who were treated at Massachusetts General Hospital and Boston Children’s Hospital. Most CDH patients (9/13) had left-sided hernia (Table 1). All four types of defect sizes (types A-D) and various comorbidities were represented in this cohort (Table 1). Except for one preterm CDH patient, all CDH patients were born term at 36-40 weeks of gestation. Age-matched BSC controls were derived from non-CDH, term (n=11) and preterm newborns (n=7) who were intubated for reasons unrelated to congenital anomalies of the lung (Table S1). CDH BSCs and non-CDH BSCs displayed no difference in cell morphology and expression of basal cell markers and NKX2.1 (Fig. 2A, compared to Fig. 1C). We also did not observe significant changes between non-CDH and CDH BSCs in proliferation rate at an average of 28-32 hours per cell cycle.

**Fig. 2.**
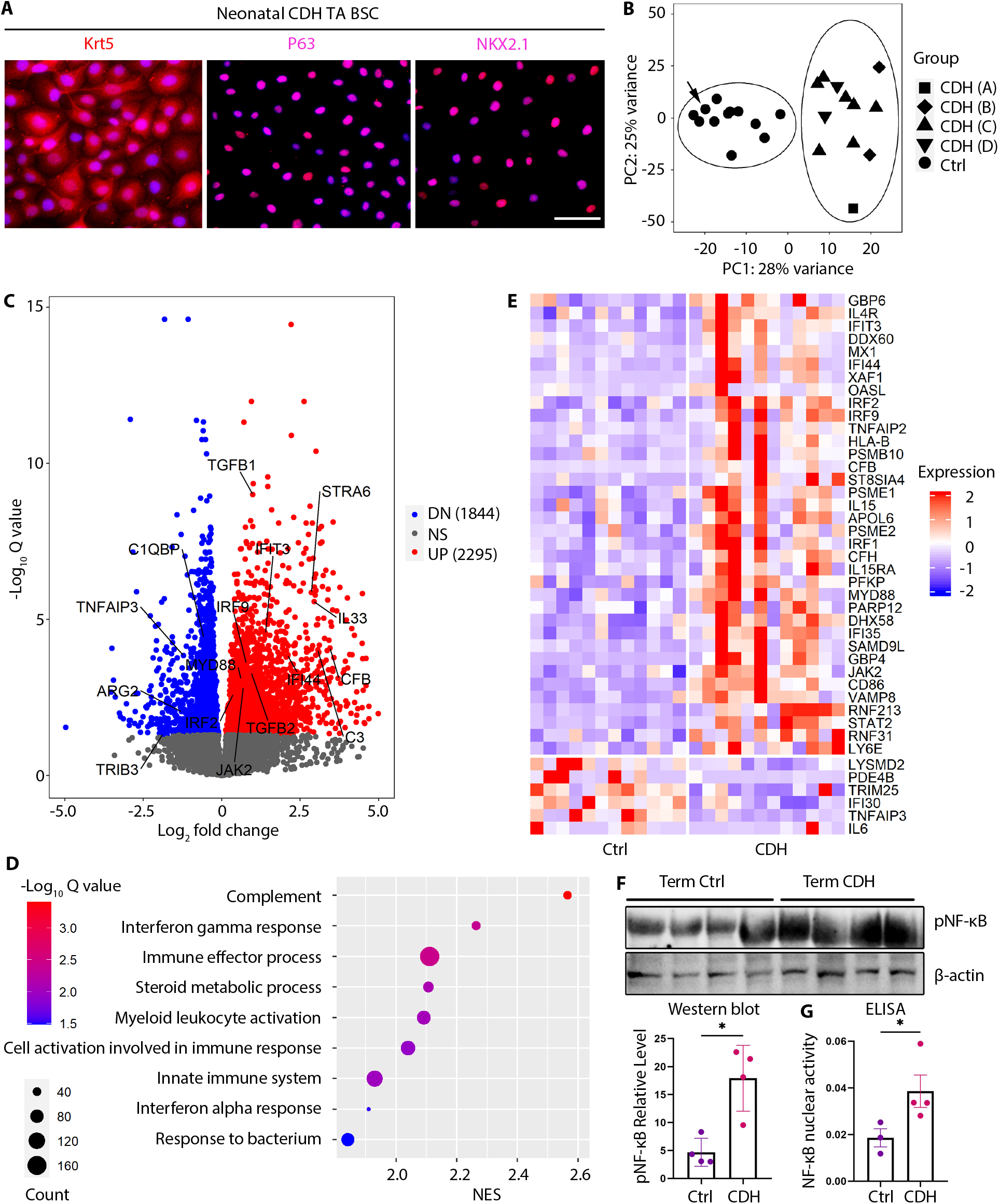
CDH BSCs have a disease-associated transcriptome and exhibit an inflammatory signature. (**A**) Representative images of antibody staining for KRT5, P63, and NKX2.1 in BSCs derived from TA of CDH newborns. Scale bar, 25μM. (**B**) Principal component analysis of bulk RNA- seq datasets of BSC samples from CDH newborns (n=12) and non-CDH, term (n=6) and preterm (n=6) newborns (Ctrl). CDH patients are labelled with A, B, C, D according to their reported type of defect size during surgical repair (62). Arrow indicates one control case with prenatal oligohydramnios associated lung hypoplasia. (**C**) Volcano plot highlighting differentially expressed genes between CDH and control BSCs with an adjusted p value <0.05. (**D**) Pathway analysis for bulk RNA-seq results of CDH and control BSCs. (**E**) Heatmap of interferon signaling pathway genes that are differentially expressed between CDH and control BSCs. (**F**) Representative Western Blot for levels of Ser536 phosphorylation of the p65 subunit of NF-κB (pNF-κB) in term, CDH (n=4) and control (n=4) BSCs. β-actin was loading control. **(G)** Activity of nuclear NF-κB measured by ELISA in CDH BSCs (n=4) and control BSCs (n=3).

To evaluate CDH-associated changes in gene expression, we performed bulk RNA-seq of the first 12 derived CDH BSC lines and 12 randomly selected control BSC lines from 6 term and 6 preterm non-CDH newborns (Table 1 and Table S1). One preterm non-CDH infant had oligohydramnios- related lung hypoplasia. Unsupervised principal component analysis (PCA) determined no difference in transcriptomic profiles of preterm and term, non-CDH BSCs (Fig. 2B). We thus combined these 12 non-CDH BSCs as one control group for further analysis. CDH BSCs derived from patients with hernia sizes ranging from type A to D clustered together and separately from controls (Fig. 2B). These findings indicate that CDH deregulates the transcriptome of BSCs in a manner distinct from prematurity and independent of the hernia size.

Compared to control BSCs, CDH BSCs had 4139 differentially expressed genes (DEGs, padj<0.05) including 1844 downregulated and 2295 upregulated genes (Fig. 2C). A disproportionally large number of DEGs in CDH were on chromosome 19 (Fig. S2A). Many DEGs in CDH BSCs were related to inflammatory responses, including *MYD88, JAK2*, interferon responsive genes that were increased, and inhibitors of the NF-κB pathway (*TNFAIP3, TRIB3*) that were reduced (Fig. 2, C and E, and Fig. S2B). Among epithelial differentiation signaling pathway genes, *STRA6* (a retinoic acid responsive gene), *TGFB1*, and *TGFB2* were significantly increased in CDH BSCs, whereas the expression of *WNT5A, WNT7B*, and *WNT9A* was similar between CDH BSCs and control BSCs (Fig. 2C and Fig. S2B).

Gene Set Enrichment Analysis (GSEA) found enrichment of DEGs in inflammatory processes including the top 2, complement and interferon gamma pathways (Fig. 2D). A heatmap of DEGs involved in the interferon gamma pathway (33) showed an overall increase in expression in CDH BSCs compared to controls (Fig. 2E). To test whether CDH BSCs had a hyperactive inflammatory response phenotype, we evaluated the status of NF-κB activation by quantifying phosphorylation of the p65 subunit. Western blot showed that CDH BSCs had a significant, ~3-fold increase in p65 phosphorylation compared to control BSCs (Fig. 2F). We also detected significantly increased DNA binding activity of nuclear NF-κB by ELISA in CDH BSCs (n=4) when compared to term control BSCs (n=3; Fig. 2G). Therefore, CDH BSCs display a proinflammatory signature.

### Changes in chromatin accessibility in CDH BSCs are associated with a proinflammatory phenotype

CDH-associated changes in gene expression were maintained in cultured BSCs after derivation from TA. Moreover, genes known to be epigenetically regulated were enriched in DEGs of CDH BSCs (Fig. 3A). To test whether epigenetic mechanisms may underlie the proinflammatory signature of CDH BSCs, we compared chromatin accessibility profiles of CDH BSCs and control BSCs by ATAC-seq (Fig. 3B). Compared to control BSCs, CDH BSCs had 14,653 peaks with increased chromatin accessibility (UP domains) and 14,518 peaks with reduced accessibility (DN domains) (Fig. 3C). We found a small but significant increase in signal density within the 1kb region surrounding the transcription start sites of genes that had at least one differentially enriched peak (Fig. 3D). *IRF2, IFIT3, MYD88, C3, CFB, TGFB1, TGFB2, and JAK2*, which were upregulated in CDH BSCs (Fig. 2, C and E, and Fig. *2B*), had increased peak signals (Fig. 3E and Fig. S2C). In contrast, downregulated genes in CDH BSCs, such as *TNFAIP3*, showed decreased peak signals (Fig. 3E). In addition, within UP and DN domains in CDH BSCs, binding motifs for transcriptional factors with well-known roles in gene activation and inflammatory responses, such as AP1 family members, NF-κB, and P63, were significantly enriched (Fig. 3F). The binding motif for epigenetic insulators, CTCF and BORIS (34), was also enriched in UP and DN domains in CDH BSCs (Fig. 3F), suggesting 3D chromatin structural domain as a potential mechanism of gene regulation. Taken together, CDH BSCs register an aberrant chromatin accessibility profile that may facilitate transcriptional deregulation of genes involved in inflammatory responses.

**Fig. 3.**
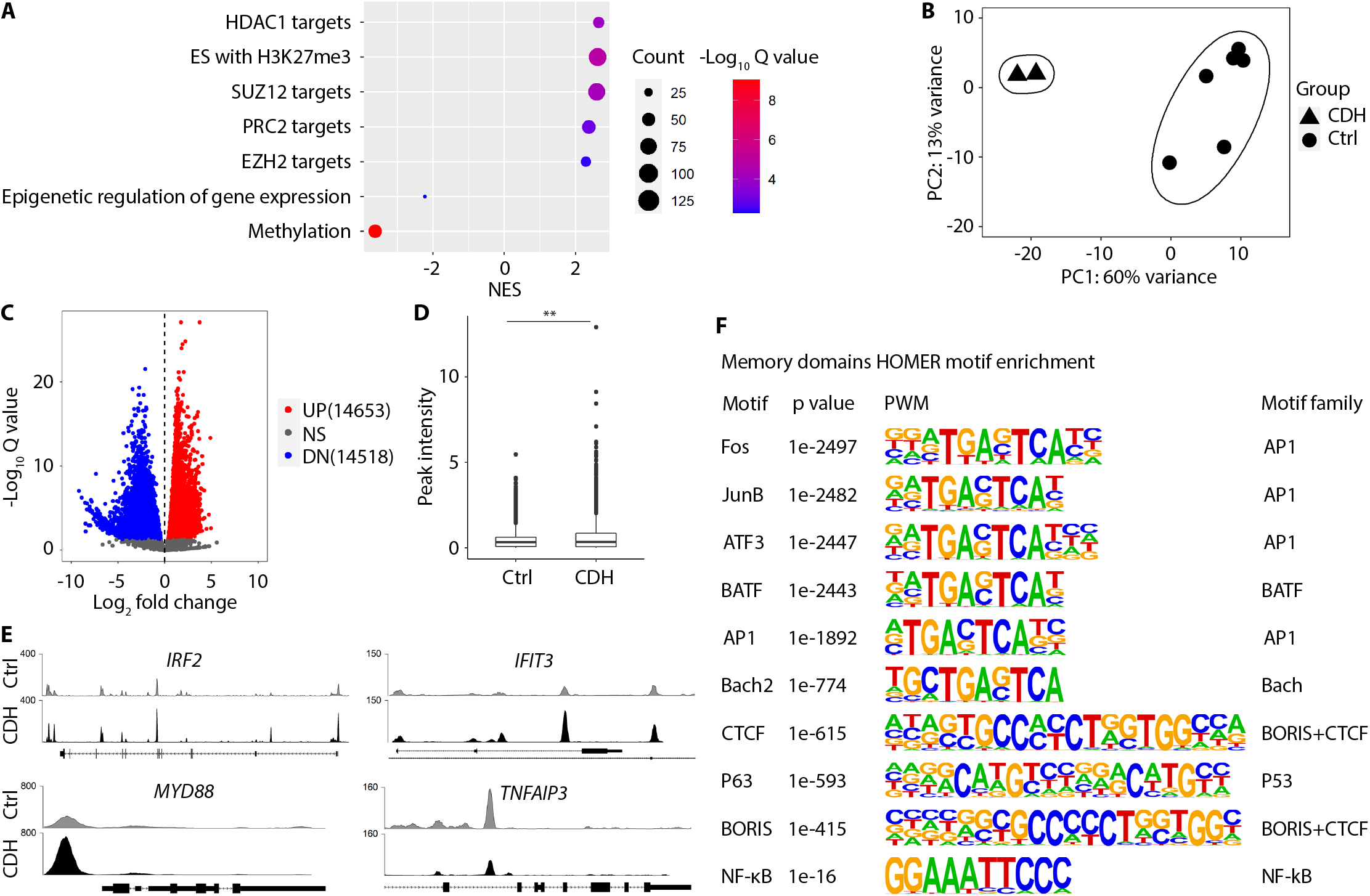
BSCs from CDH newborns exhibit changes in chromatin accessibility. (**A**) Enrichment of target genes of epigenetic regulation among differentially expressed genes (DEGs) by bulk RNA- seq between CDH and control BSCs. UP, upregulated; DN, downregulated. (**B**) Principal component analysis of ATAC-seq datasets comparing CDH BSCs (n=2) and control BSCs (n=6). (**C**) Volcano plot highlighting significantly altered chromatin accessibility between CDH BSCs and control BSCs. (**D**) Summary of peak intensity within 1 KB surrounding the transcription start site of genes with at least one enriched peak between CDH and control BSCs. (**E**) Detailed examination of changes in peak signals within regulatory sequences of select genes. (**F**) HOMER motif analysis of chromatin accessibility. Students t-test was used for statistical analysis, ** p<0.01.

### CDH BSCs exhibit heterogenous defects in epithelial differentiation

We tested whether CDH BSCs had changes in epithelial differentiation using day 21 ALI cultures (Fig. 4A). All 16 control BSC lines, including 5 preterm and 11 term lines, generated stratified airway epithelium with no difference in the abundance of residual BSCs (KRT5^+^ cells, 63.3%±4.0% (preterm) vs 56.3%±4.5% (term), p=0.44; P63^+^ cells, 28.8%±1.7% (preterm) vs 34.1%±2.9% (term), p=0.25), CC10^+^ club cells (4.2%±1.6% (preterm) vs 3.5%±0.9% (term), p=0.74), Muc5AC^+^ goblet cells (2.0%±0.7% (preterm) vs 3.8±1.2% (term), p=0.66), and RFX3^+^ ciliated cells (31.0±5.4% (preterm) vs 34.4%±6.2% (term), p=0.91) (Fig. 4, B and C). These results indicate that gestational age at birth has no effect on the differentiation potential of BSCs, which is consistent with our findings that preterm and term, non-CDH BSCs have similar transcriptomic profiles (Fig. 2B).

**Fig. 4.**
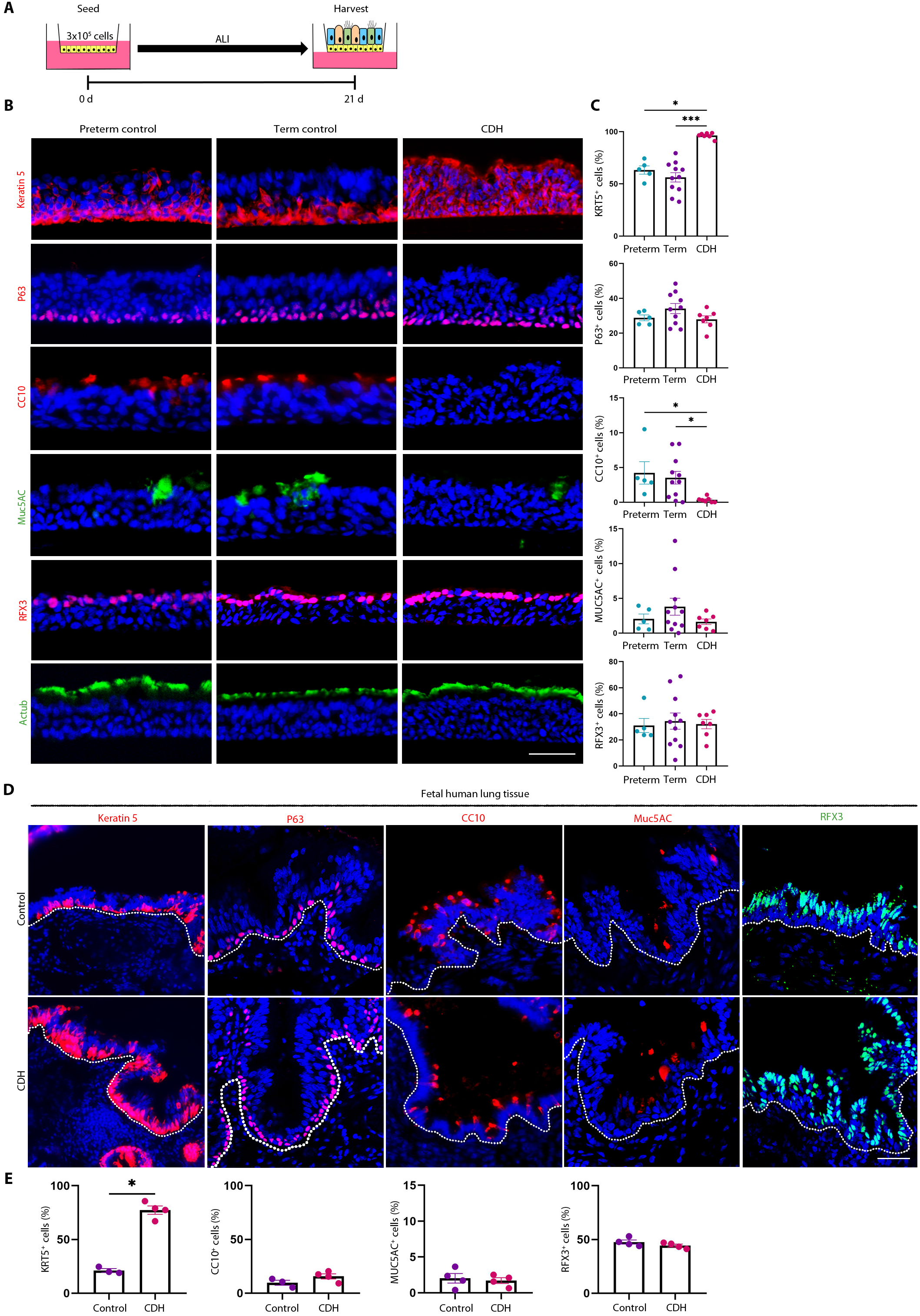
CDH BSCs have epithelial differentiation defects in ALI culture and *in vivo*. (**A**) Schematic of ALI culture of preterm, term, and CDH BSCs for 21 days. (**B**) Representative fluorescence images of staining for markers of BSCs (KRT5 and P63), Club cells (CC10), goblet cells (Muc5AC), and ciliated cells (RFX3 and acetylated tubulin (Actub)) using cross sections of fixed ALI cultures. Scale bar, 25 μM. (**C**) Quantification of the relative abundance of different epithelial cell types based on the staining of ALI culture in triplicates. More than 6000 nuclei in five 20X images were counted for each marker for each ALI culture. Each dot represents one BSC line. (**D & E**) Representative fluorescence images and quantification of immunostaining for KRT5, P63, CC10, MUC5AC and RFX3 and in term, control (n=3) and CDH (n=4) fetal lung sections. Scale bar, 50 μM. More than 2,000 nuclei in 5 images were counted per sample. Bar graphs represent mean±SEM. Statistical analyses were performed using Kruskall-Wallis test followed by a post-hoc corrected Dunn’s test for comparisons of multiple experimental groups and Mann–Whitney U test for comparisons between two groups. Significance indicated as *p<0.05, **p<0.01, and ***p<0.001.

In contrast, 6 out of 13 CDH cell lines failed to differentiate in ALI after multiple attempts at varied cell seeding densities. The other 7 CDH lines were able to differentiate; however, the generated airway epithelium was abnormal. KRT5 was retained in nearly 100% of cells (96.3%±1.0% (CDH), p<0.05) including cells already expressing differentiated epithelial markers. Moreover, club cell numbers were significantly reduced (0.34%±0.1% (CDH), p<0.05)) compared to control ALI cultures (Fig. 4, B and C, and Fig. S3) (35). CDH BSCs had no significant change in differentiation into ciliated cells and goblet cells (Fig. 4, B and C). Furthermore, the differentiation phenotypes of CDH BSCs were not caused by delayed differentiation, as they remained unchanged even after prolonged ALI culture for 35 days (Fig. S4). Lastly, only 105 genes were differentially expressed between the 7 CDH BSCs that were able to differentiate and 6 CDH BSCs that failed to differentiate (Fig. S5), which precludes reliable pathway predictions for further testing. Taken together, CDH BSCs were defective in differentiation; however, their differentiation phenotypes are not identical despite a shared proinflammatory signature.

### CDH BSCs partially reproduce the epithelial phenotype in CDH patients

To assess epithelial defects in CDH patients, we examined lung sections from fetopsy samples of term, CDH (n=4) and non-CDH fetuses (n=3) that were not treated pre- or post-natally. First, the non-CDH fetal airway epithelium *in vivo* contained differentiated epithelial cell types in relative abundance similar to that in control ALI cultures. Second, CDH lungs had more abundant KRT5^+^ cells than non-CDH controls (70.1%±5.1% vs. 30.4%±5.9%, p<0.01) (Fig. 4, D and E), which is consistent with altered epithelial differentiation *in vitro* (Fig. 4, B and C). We did not observe changes in the abundance of p63^+^ basal cells, RFX3^+^ ciliated cells and MUC5AC^+^ goblet cells in CDH airways compared to controls (Fig. 4, D and E). However, we found a significant number of ciliated cells marked by Actub that were also KRT5^+^ in CDH airways, which resembles our observations in CDH ALI cultures (Fig S3, C and D). We found no change in CC10^+^ club cells in intrapulmonary airway epithelium of CDH fetuses (Fig. 4, D and E). The discrepancy in club cell differentiation between *in vitro* and *in vivo* assays may be explained by a compensatory increase in club cell differentiation from other epithelial progenitors that co-exist with BSCs in human conducting airways (36, 37).

### The proinflammatory signature of CDH BSCs underlies abnormal epithelial differentiation

To test whether the proinflammatory signature of CDH BSCs may deregulate epithelial differentiation, we treated CDH BSCs with dexamethasone (DXM), an anti-inflammatory drug approved for induction of lung maturation in cases of preterm birth (38). We first tested whether DXM could restore epithelial differentiation in the subset of CDH BSCs that were able to differentiate (Fig. 5A and Fig. S6). We showed that epithelial differentiation of CDH BSCs was normalized by minimally two weeks of DXM treatment (10 μM), including 1 week before ALI and during the first week in ALI (Fig. 5, A-C, and Fig. S6). This was documented by reduced KRT5^+^ cells (96.3%±1.0% vs 39.9%±1.9%, p<0.0001) and rescued club cell differentiation (CC10^+^, 0.34%±0.1% vs 9.0%±2.0%, p<0.01). Other cell types in ALI culture were unchanged by DXM treatment (Fig. 5, B and C). Accordingly, DXM treatment reduced the level of p65 phosphorylation in CDH BSCs by approximately 50% (Fig. 5D). These beneficial effects of DXM were not improved by prolonged treatment or at a higher concentration (10 mM, Fig. S6). DXM treatment had no effect on epithelial differentiation of term, control BSCs (Fig. S7). Furthermore, DXM treatment failed to rescue the other 6 CDH BSC lines that were incapable of differentiation in ALI. Additionally, a specific NF-κB inhibitor had similar rescue effects as DXM on CDH BSC differentiation, without any effect on controls (Fig. S8, B-E). Lastly, since the IL-6/STAT3 pathway plays established roles in inflammation and BSC differentiation (39), we evaluated whether STAT3 activation may be involved in the differentiation phenotypes of CDH BSCs. We found no difference in baseline levels of phosphorylated, active STAT3 in control and CDH BSCs (Fig. S9A). Correspondingly, blockade of STAT3 activity had no effect on the differentiation of CDH BSCs (Fig. S9, B-D).

**Fig. 5.**
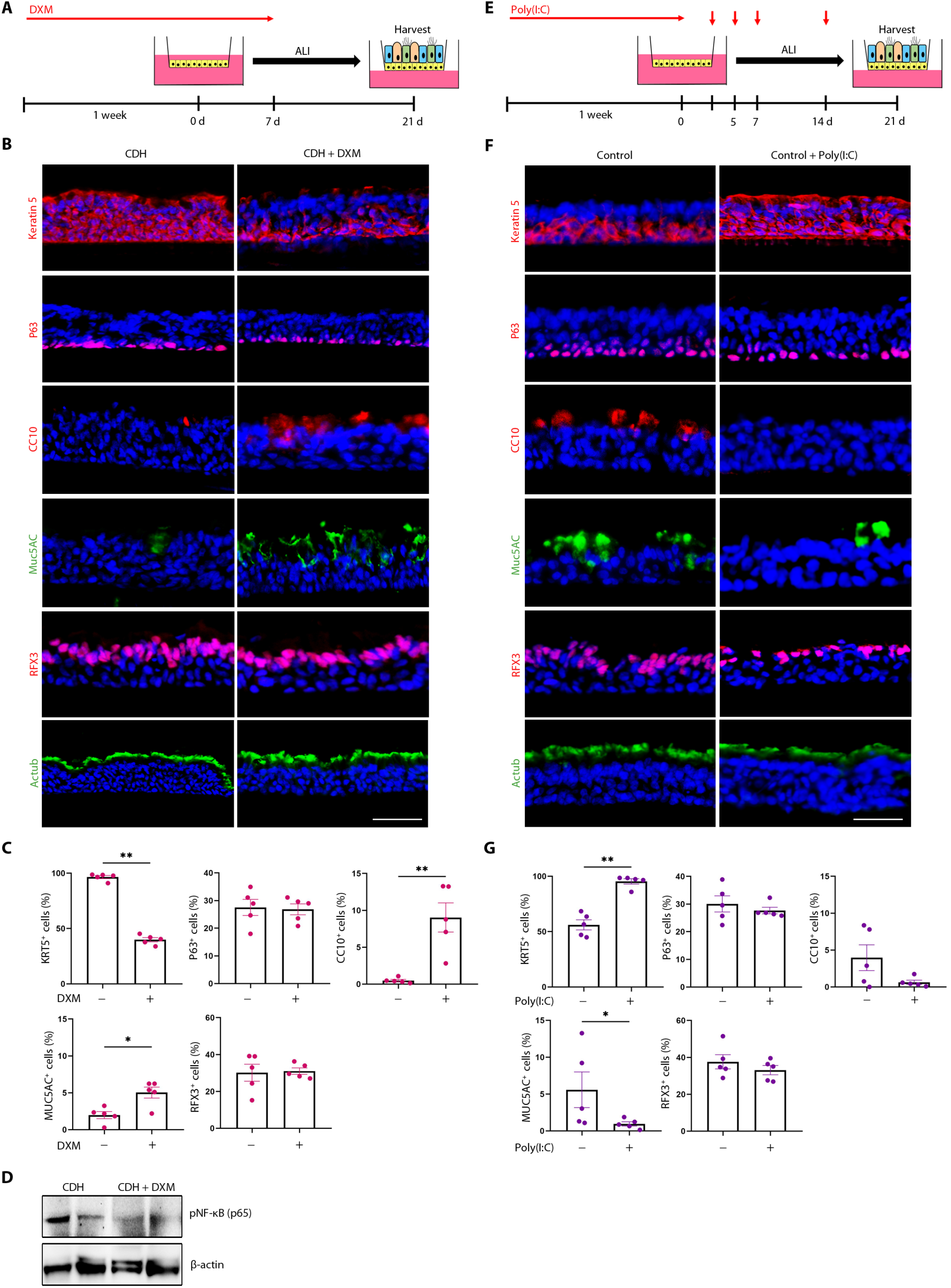
Dexamethasone rescues differentiation defects of CDH BSCs in ALI cultures while poly(I:C) stimulation of control BSCs induces differentiation defects similar to those of CDH BSCs. (**A**) Schematic of DXM (10 μM) treatment of CDH BSCs before and during ALI culture. (**B**) Representative fluorescence images of antibody staining for epithelial cell markers in day 21 ALI cultures of CDH BSCs (n=5 lines) with and without DXM treatment. (**C**) Quantification of each labelled cell types in ALI cultures. (**D**) Representative Western Blot for phosphorylated Ser536 in the p65 subunit (pNF-κB) in CDH BSCs with and without DXM treatment for 1 week during cell culture. β-actin was loading control. (**E**) Schematic of Poly (I:C) (10 μM) treatment of control BSCs (n=3 lines) before and during ALI culture. (**F**) Representative fluorescence images of antibody staining for epithelial cell markers in day 21 ALI cultures of control BSCs with and without poly(I:C) treatment. The relative abundance of each labelled cell type was quantified in (**G**). ALI culture was performed in triplicates and more than 6000 cells were counted using stained sections of fixed ALI cultures for each BSC line. *p<0.05 and **p<0.01 by Mann–Whitney U test. Scale bar, 25 μM.

Poly(I:C), a mimic of double-stranded viral RNA, induces activation of the NF-κB and interferon pathways in multiple cell types, including airway epithelial cells (40, 41). Since CDH BSCs exhibited NF-κB hyperactivity and elevated levels of interferon response gene expression (Fig. 2, D-F), we tested whether poly(I:C) treatment of control BSCs mimic the impact of CDH on epithelial differentiation. To do so, control BSCs (n=5) were treated with 10 μM Poly(I:C) for 1 week before ALI and on days 3, 5, 7, and 14 during ALI culture (Fig. 5E). Poly (I:C) treated cell lines responded with elevated NF-κB nuclear transcriptional activity (Fig. S8F). Day 21 ALI cultures of poly(I:C)- treated control BSCs showed an increase in KRT5^+^ cells to almost 100% and a reduction in the number of club and goblet cells (Fig. 5, F and G), which is similar to CDH BSC differentiation (Fig. 4, B and C). In conclusion, the results of DXM and NF-κB inhibitor rescue assays in CDH BSCs and poly (I:C) disease mimicking assays in control BSCs indicate that the proinflammatory signature in CDH BSCs, manifested by hyperactive NF-κB and interferon pathways, is functionally linked to abnormal epithelial differentiation in a subset of CDH BSCs.

### DXM treatment can rescue the tracheal epithelial differentiation phenotype in the rat CDH model

To determine whether the beneficial effects of DXM on CDH BSC-based *in vitro* model could be reproduced *in vivo*, DXM was administrated to the nitrofen rat CDH model between E10.5 - E12.5 (Fig. 6A) when BSCs are generated in the tracheal airway epithelium based on lineage-tracing in mice (24). Nitrofen exposure (E9) induced diaphragmatic hernia in 53% fetuses (16/30) and caused lung hypoplasia in 100% of fetuses (30/30) at E21.5 in our study. Consistent with previous findings (42, 43), we found that DXM treatment improved both lung size (Fig. S10) and average lung weight (control (0.21g), nitrofen no hernia (0.09g), nitrofen with hernia (0.07g), nitrofen+DXM (0.15g)) and reduced the occurrence of diaphragmatic hernia (3/19). Nitrofen exposure significantly increased the number of KRT5^+^ epithelial cells and reduced club cell numbers in tracheal epithelium (Fig. 6, B and C), which resemble epithelial abnormalities in ALI cultures of CDH BSCs and the increased KRT5^+^ cells in fetal CDH lungs (Fig. 4). In contrast, club cells in intrapulmonary airways of nitrofen- exposed fetuses appeared unaffected (Fig. 6D). Of note, club cells in rodent intrapulmonary airways are generated from epithelial progenitors distinct from BSCs (44). These findings suggest that BSCs may be selectively affected in the nitrofen model. Following DXM treatment, the tracheal epithelium in nitrofen-exposed fetuses was normalized as evidenced by KRT5 localization to the basal cell layer and recovered club cell differentiation (Fig. 6, B and C). DXM treatment also restored air space in alveoli of nitrofen-exposed fetuses (Fig. 6, E and F). Therefore, early embryonic administration of steroid effectively targets lung phenotypes in the rat nitrofen model of CDH.

**Fig. 6.**
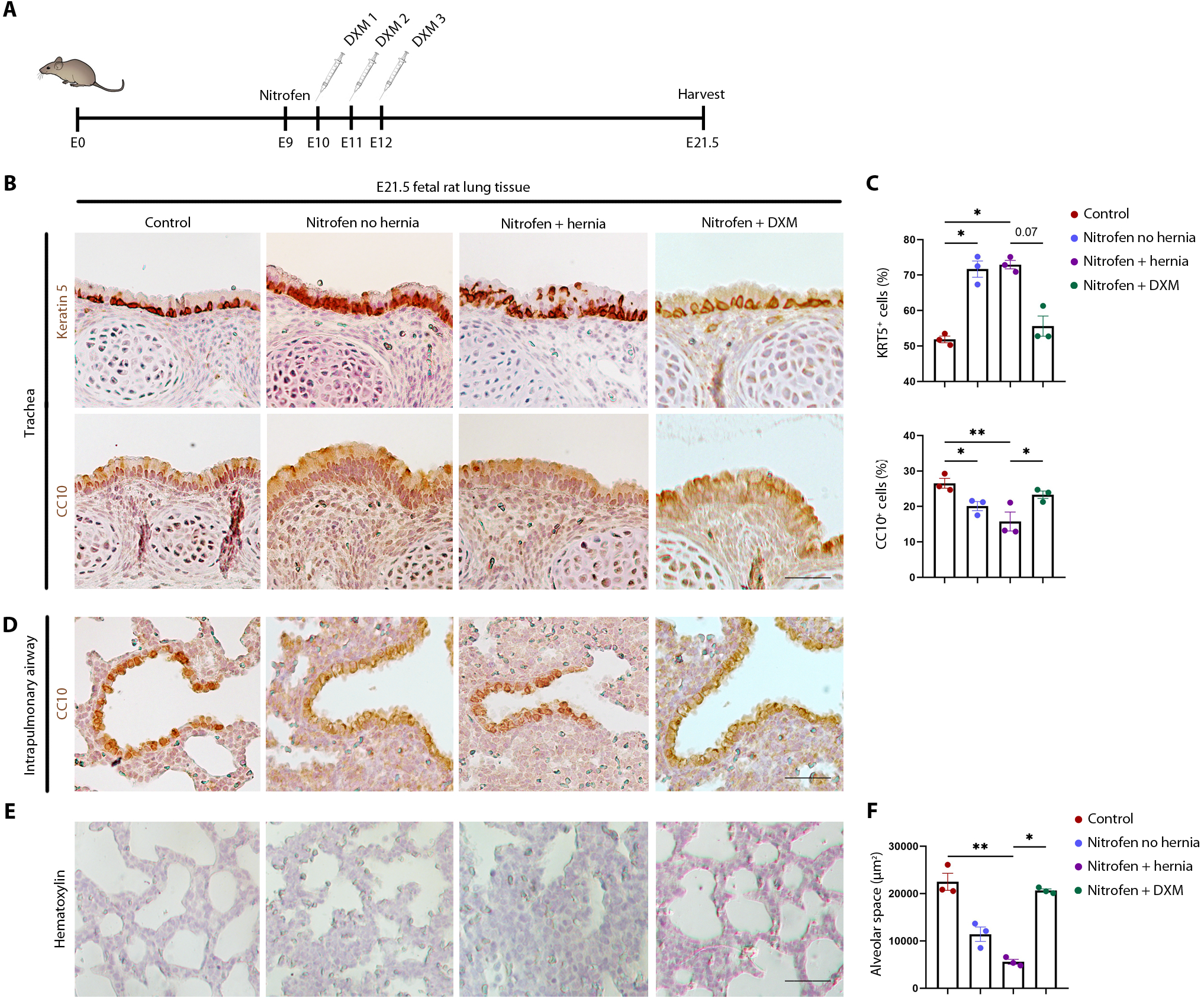
DXM treatment restores normal tracheal epithelium in rat fetuses of the nitrofen model of CDH. (**A**) Schematic of nitrofen (100mg in olive oil) gavage at E9.5 and DXM treatment (0.25 μg/g at E10.5 and E11.5 and 0.125 μg/g at E12.5). Control dams received olive oil. Fetuses were analyzed at E21.5. (**B**) Representative images of antibody staining for KRT5 and CC10 in rat fetal tracheas of controls (n=3), nitrofen with no hernia (n=4), nitrofen with hernia (n=4), and nitrofen with DXM treatment (n=3). (**C**) Quantification of the relative abundance of KRT5^+^ and CC10^+^ cells in tracheal epithelium of each group. (**D**) Representative CC10 staining of intrapulmonary airway sections of each group. (**E**) Representative hematoxylin staining of distal lung sections of each group. (**F**) Quantification of air space in the alveoli of each group. For quantification, more than three 20X images for each sample were quantified. Each dot represents one sample. Bar graphs show mean±SEM. *p<0.05; **p<0.01 by Kruskall-Wallis test followed by a post-hoc uncorrected Dunn’s test.

## DISCUSSION

Here, we present a robust methodology of BSC derivation from TA to model epithelial defects of CDH newborns *in vitro*. We provide evidence that CDH BSCs *in vitro* reproduce epithelial defects in human fetal CDH lungs and tracheas of the nitrofen rat model of CDH. This translational approach enables the identification of a developmental defect of BSCs in CDH manifested by a proinflammatory signature that disrupts epithelial differentiation. Our approach of TA BSC derivation can now be expanded to other neonatal lung diseases for disease modelling and therapeutic testing *in vitro*. The proinflammatory signature in CDH BSCs is in accordance with a recent proteomic study that found elevated inflammatory responses in nitrofen CDH lungs (45). Specifically, elevated NF-κB in proximal epithelial cells during murine lung development can cause distal lung hypoplasia (46). Additional studies emphasize the crosstalk between inflammatory mediators and pathways involved in branching morphogenesis and epithelial differentiation (47–52). By expanding on these findings, our results may inform pathogenesis of lung defects in CDH.

We demonstrate that TA are an alternative source of patient-specific lung cells to circumvent technical difficulties of accessing fresh lung tissue in newborns. TA-derived BSCs also offer additional advantages, including time- and cost-effectiveness, compared to generating patient specific human lung cells using induced pluripotent stem cell-based (iPSC) approaches (53, 54). More importantly, by maintaining both the genetic and epigenetic landscapes, TA BSCs enable the characterization of patient-specific phenotypes with greater fidelity than iPSC-derived lung progenitors which have undergone epigenetic reprogramming (55).

We acknowledge that our study using one type of epithelial progenitors harvested around birth is limited in reproducing complex interactions between multiple cell types during all stages of lung development. However, this concern is mitigated by similar epithelial phenotypes observed in our BSC culture system, CDH patient samples, and the nitrofen rat model of CDH. Future studies will investigate how BSC differentiation phenotypes and abnormal programming relate to lung hypoplasia seen in CDH patients.

While previous studies suggested that CDH lungs may be developmentally premature (1, 2), we show that CDH BSCs display divergent transcriptome, chromatin accessibility and differentiation when compared to preterm BSCs. Thus, CDH BSCs are not simply in a premature state. Furthermore, the size of the diaphragmatic hernia has no significant impact on gene expression or differentiation capacity of CDH BSCs. These findings suggest the proinflammatory signature of CDH BSCs as an intrinsic mechanism of abnormal epithelial differentiation in CDH.

Currently no prenatal medical treatment can effectively improve lung function in CDH patients. Our findings of beneficial effects of antenatal dexamethasone on conducting airway epithelium is in line with improved alveolar morphology in the nitrofen rat model and surgical sheep model of CDH following steroid treatment (42, 43, 56). However, a small clinical trial assessing the effect of betamethasone during last trimester CDH pregnancies did not show improved outcomes (57). The caveats of this trial include the small number of patients, dosage and timepoint of treatment. Recent reports show encouraging results for the prenatal treatment of severe CDH cases using fetoscopic endoluminal tracheal occlusion (FETO) (58). Novel therapeutics, such as amniotic fluid derived extracellular vesicles can promote prenatal lung maturation in CDH animal models (59, 60). Future studies could explore how such therapeutics can rescue CDH phenotypes in our *in vitro* model and large animal models before clinical testing in CDH patients when applied alone or in combination with FETO (61). Lastly, TA BSCs derived from larger CDH patient cohorts, in combination with genetic mapping of patient-specific de novo mutations, is warranted to deepen our understanding of the pathogenesis of lung defects in CDH.

## ACKNOWLEDGEMENT

We would like to thank Jennifer Lyu and Gheed Murtadi for coordinating TA collection from CDH newborns. This work is supported by a grant from the German Research Foundation (DFG) to RW (461188606); NIH grants to PKD (NICHD 2PO1HD068250) and PHL (R21A1156597); and funds from the Department of Pediatrics at MGH for the Lung Cell Bank to XA.

