## Supplemental Material for "A tracheal aspirate-derived airway basal cell model reveals a proinflammatory epithelial defect in congenital diaphragmatic hernia"

### SUPPLEMENTARY MATERIALS

**Supplementary Table 1:** Demographic and clinical information of preterm and term patients from whom TA BSCs were derived for this study.

| Patient Study ID | Sex | Survival (Y/N) | Gestational Age (weeks) | Reason for Intubation | Differentiated (Y/N) | Assays Performed |
|---|---|---|---|---|---|---|
| 1 | M | Y | 39 | NAS | Y | R, A, D |
| 2 | F | N | 41 | HIE | Y | R, A, D |
| 3 | F | Y | 38 | Fetal maternal hemorrhage, PPHN | Y | R, A, D |
| 4 | M | Y | 41 | MAS, PPHN, HIE | Y | R, D |
| 5 | M | Y | 40 | Pneumothorax | Y | R, D |
| 6 | F | Y | 37 | HIE, MAS, PPHN | Y | R, D |
| 7 | M | Y | 41 | Amniotic fluid aspiration, PNA | Y | D |
| 8 | F | Y | 39 | MAS, PPHN | Y | D |
| 9 | M | Y | 39 | Severe HIE, PPHN | Y | D |
| 10 | M | Y | 40 | MAS, PPHN, pulmonary hemorrhage, | Y | D |
| 11 | M | Y | 39 | Severe HIE, PPHN | Y | D |
| 12 | F | Y | 24 | RDS | Y | R, A, D |
| 13 | F | Y | 27 | PET | Y | R, A, D |
| 14 | F | N | 24 | Strep anginosus | Y | R, A, D |
| 15 | F | Y | 27 | Abruption | Y | R, D |
| 16 | M | Y | 25 | Cong candida | Y | D |
| 17 | F | Y | 26 | rBMZ, PET | Not Assayed | R |
| 18 | F | Y | 27 | DCDA twin | Not Assayed | R |

***NAS:** neonatal abstinence syndrome, **HIE:** hypoxic-ischemic encephalopathy, **PPHN:** persistent pulmonary hypertension of the newborn, **AVM:** arteriovenous malformations, **PNA:** pulmonary nodular amyloidosis, **MAS:** macrophage activation syndrome, **RDS:** respiratory distress syndrome, Pre-eclampsia, **BMZ:** betamethasone, **HELLP:** hemolysis, elevated liver enzymes, and low platelets, **DCDA:** dichorionic diamniotic

***R** = RNA Sequencing, **A** = ATAC Sequencing, **D** = Differentiation

**Supplementary Table 2:** Primer sequences used in this study.

| Target | Primer sequence |
|---|---|
| Human ACTB-F | 5’-CACCATTGGCAATGAGCGGTTC-3’ |
| Human ACTB-R | 5’-AGGTCTTTGCGGATGTCCACGT-3’ |
| Human IL33-F | 5’-GCCTGTCAACAGCAGTCTACTG-3’ |
| Human IL33-R | 5’-TGTGCTTAGAGAAGCAAGATACTC-3’ |
| Human IL1RL2-F | 5’-TGTGCTTAGAGAAGCAAGATACTC-3’ |
| Human IL1RL2-R | 5’-GGACCACAATGACAATCAGCCTC-3’ |
| Human ANKLE1-F | 5’-TCCGAGCACTTGGTGAGAATCC-3’ |
| Human ANKLE1-R | 5’-CAGTTCTAGGCTGTGCCCTGAA-3’ |
| Human ALDH3A1-F | 5’-CTCGTCATTGGCACCTGGAACT-3’ |
| Human ALDH3A1-R | 5’-CTCGCCATGTTCTCACTCAGCT-3’ |
| Human MYD88-F | 5’-GAGGCTGAGAAGCCTTTACAGG-3’ |
| Human MYD88-R | 5’-GCAGATGAAGGCATCGAAACGC-3’ |
| Human JAK2-F | 5’-CCAGATGGAAACTGTTCGCTCAG-3’ |
| Human JAK2-R | 5’-GAGGTTGGTACATCAGAAACACC-3’ |
| Human STRA6-F | 5’-CTGGAAAGCCTTCACTGAGGAG-3’ |
| Human STRA6-R | 5’-CTGATCTGCCAGAGTCTCGAAG-3’ |
| Human TGFB1-F | 5’-TACCTGAACCCGTGTTGCTCTC-3’ |
| Human TGFB1-R | 5’-GTTGCTGAGGTATCGCCAGGAA-3’ |
| Human TGFB2-F | 5’-AAGAAGCGTGCTTTGGATGCGG-3’ |
| Human TGFB2-R | 5’-ATGCTCCAGCACAGAAGTTGGC-3’ |
| Human TGFBR2-F | 5’-GTCTGTGGATGACCTGGCTAAC-3’ |
| Human TGFBR2-R | 5’-GACATCGGTCTGCTTGAAGGAC-3’ |
| Human WNT5A-F | 5’-TACGAGAGTGCTCGCATCCTCA-3’ |
| Human WNT5A-R | 5’-TGTCTTCAGGCTACATGAGCCG-3’ |
| Human WNT7B-F | 5’-AGAAGACCGTCTTCGGGCAAGA-3’ |
| Human WNT7B-R | 5’-AGTTGCTCAGGTTCCCTTGGCT-3’ |
| Human WNT9A-F | 5’-AGTGCCAGTTCCAGTTCCGCTT-3’ |
| Human WNT9A-R | 5’-AGGAGATGGCATAGAGGAAGGC-3’ |
| Human ZFP82-F | 5’-AATACCTGGACTTGGAACAAAAGG-3’ |
| Human ZFP82-R | 5’-TCCTCACAACTTTCCAAGGCTCT-3’ |
| Human ZNF844-F | 5’-GACCAGAACATTGAAGATCAGTAC-3’ |
| Human ZNF844-R | 5’-CAGTGTGTCATCTGGAATCTGGC-3’ |

**Supplementary Table 3:** Primary Antibodies used in this study.

| Antigen | Catalog # | Company | Concentration | | | |
|---|---|---|---|---|---|---|
| | | | BSCs and ALI sections | Western Blot | Human Tissue | Rat Tissue |
| KRT5 | ab53121 | Abcam (Cambridge, UK) | 1:100 | - | 1:100 | 1:200 |
| p63 | 13109S | Cell Signaling Technology (Danvers, MA) | 1:100 | - | 1:100 | - |
| NKX2.1 | sc-13040 | Santa Cruz Biotechnology (Dallas, TX) | 1:100 | - | - | - |
| CC10 | HPA031828 | Sigma-Aldrich (St. Louis, MO) | 1:100 | - | 1:50 | - |
| MUC5AC | MA5-12178 | Thermo Scientific (Waltham, MA) | 1:100 | - | 1:100 | - |
| RFX3 | HPA035689 | Sigma-Aldrich (St. Louis, MO) | 1:100 | - | 1:100 | - |
| Anti-tubulin | T6793 | Sigma-Aldrich (St. Louis, MO) | 1:100 | - | 1:100 | - |
| CC10 | sc-9772 | Santa Cruz Biotechnology (Dallas, TX) | - | - | - | 1:100 |
| pNF-κB | 3033T | Cell Signaling Technology (Danvers, MA) | - | 1:1000 | - | - |
| pSTAT3 | 9145S | Cell Signaling Technology (Danvers, MA) | - | 1:1000 | - | - |
| β-actin | A5441 | Sigma-Aldrich (St. Louis, MO) | - | 1:2000 | - | - |

KRT5: marker of basal cells
p63: marker of basal cells
CC10: marker of club cells
MUC5AC: marker of goblet cells
RFX3: marker of ciliated cells
Anti-tubulin: marker of ciliated cells

**Table S4.** Secondary Antibodies used in this study.

| Antibody | Conjugate | Catalog # | Company | Concentration | | | |
|---|---|---|---|---|---|---|---|
| | | | | BSCs and ALI sections | Western Blot | Human Tissue | Rat Tissue |
| Goat anti-rabbit IgG | Alexa Flour 594 | A11037 | Thermo Scientific (Waltham, MA) | 1:200 | - | 1:200 | - |
| Goat anti-mouse IgG | Alexa Flour 488 | A11029 | Thermo Scientific (Waltham, MA) | 1:200 | - | 1:200 | - |
| Goat anti-rabbit IgG | biotin | BA-1000 | Vector Laboratories (Newark, CA) | - | - | - | 1:200 |
| Goat anti-mouse IgG | biotin | BA-9200 | Vector Laboratories (Newark, CA) | - | - | - | 1:250 |
| Goat anti-rabbit IgG | HRP | 7074S | Cell Signaling Technology (Danvers, MA) | - | 1:2000 | - | - |
| Goat anti-mouse IgG | HRP | G-21040 | Thermo Scientific (Waltham, MA) | - | 1:2000 | - | - |

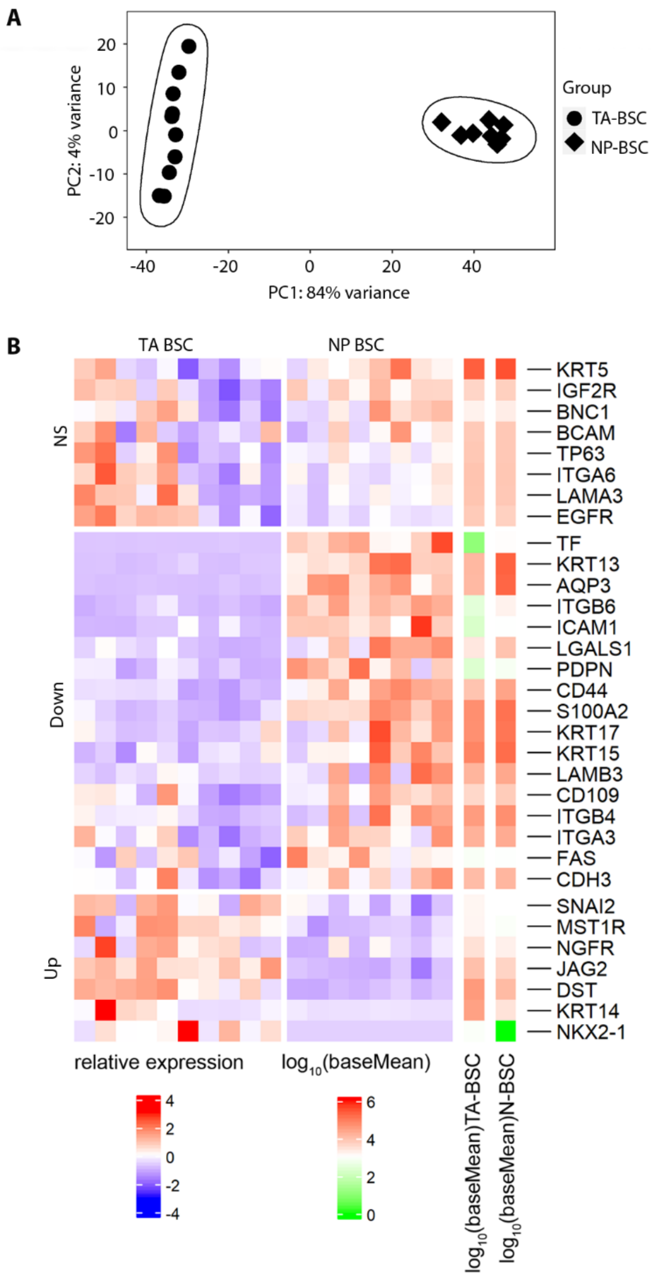


**Fig. S1. TA BSCs have a transcriptome profile distinct from nasopharyngeal (NP) BSCs. (A)** Principal component analysis of bulk RNA-seq results of neonatal TA BSCs (n=10 lines) and an already published dataset (Shui et al.[1]) of neonatal NP BSCs (n=8 lines). **(B)** Heatmap of basal stem cell marker gene expression in TA BSCs and NP BSCs. NS, not significant. Down, genes significantly downregulated in TA BSCs compared to NP BSCs. Up, genes significantly upregulated in TA BSCs compared to NP BSCs. Genes with an adjusted p value <0.05 are considered as significantly changed.

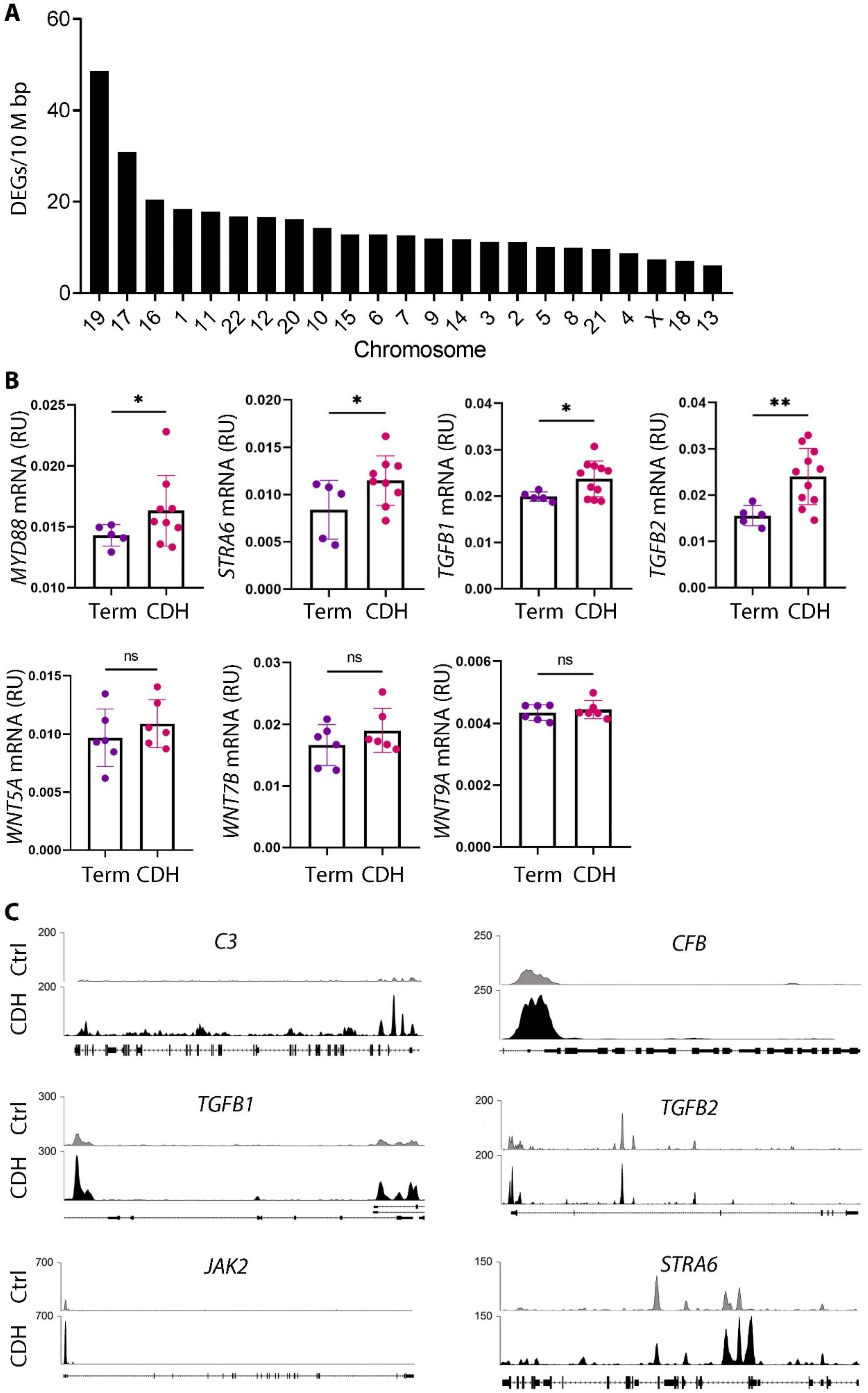


**Fig. S2. CDH BSCs exhibit significant changes in gene expression and chromatin accessibility compared to non-CDH, control BSCs.** (**A**) The relative distribution of differentially expressed genes (DEGs) on each chromosome. (**B**) Gene expression of *MYD88*, *STRA6*, *TGFB1*, *TGFB2, WNT5A, WNT7B, and WNT9A* in CDH BSCs compared to term control BSCs by RT-qPCR. Each dot represents one BSC line. The assay was performed in duplicates for each sample. *p<0.05 and **p<0.01 by Mann-Whitney U test. (**C**) Peaks of chromatin accessibility for select genes including *C3*, *CFB*, *TGFB1*, *TGFB2*, *JAK2*, and *STRA6*.

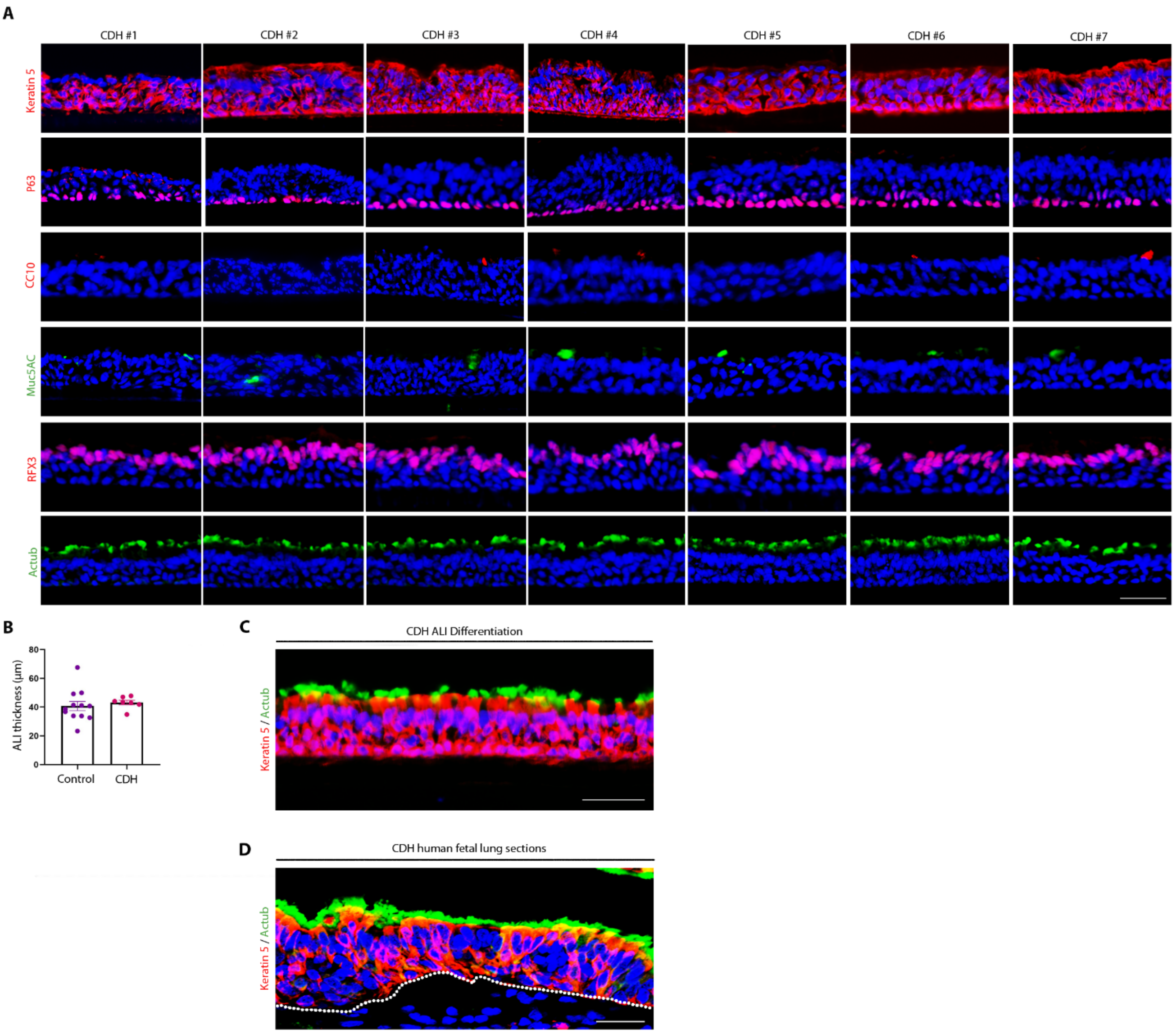


**Fig. S3. Seven CDH BSC lines were able to differentiate into various epithelial cell types in ALI.** (**A**) Representative fluorescence images of epithelial marker staining in day 21 ALI culture of the 7 CDH BSC lines that were able to differentiate. Sections were stained for markers of basal stem cells (KRT5 and P63), club cells (CC10), goblet cells (Muc5AC), and ciliated cells (RFX3 and acetylated tubulin (Actub)). Nuclei were stained by DAPI. (**B**) Quantification of the thickness of the epithelial layer in day 21 ALI culture of CDH BSCs and non-CDH, preterm and term BSCs. Each dot represents one BSC line. Bar graph represents mean ± SEM for each experimental group. At least 3 technical replicates for each line were evaluated. Scale bar, 25 µm. **(C)** Representative fluorescence images of co-staining for Keratin 5 and acetylated tubulin (Actub) in cross sections of fixed ALI cultures. Scale bar, 25 µm. **(D)** Representative fluorescence images of co-staining for Keratin 5 and acetylated tubulin (Actub) in human fetal CDH lungs. Scale bar, 50 µm.

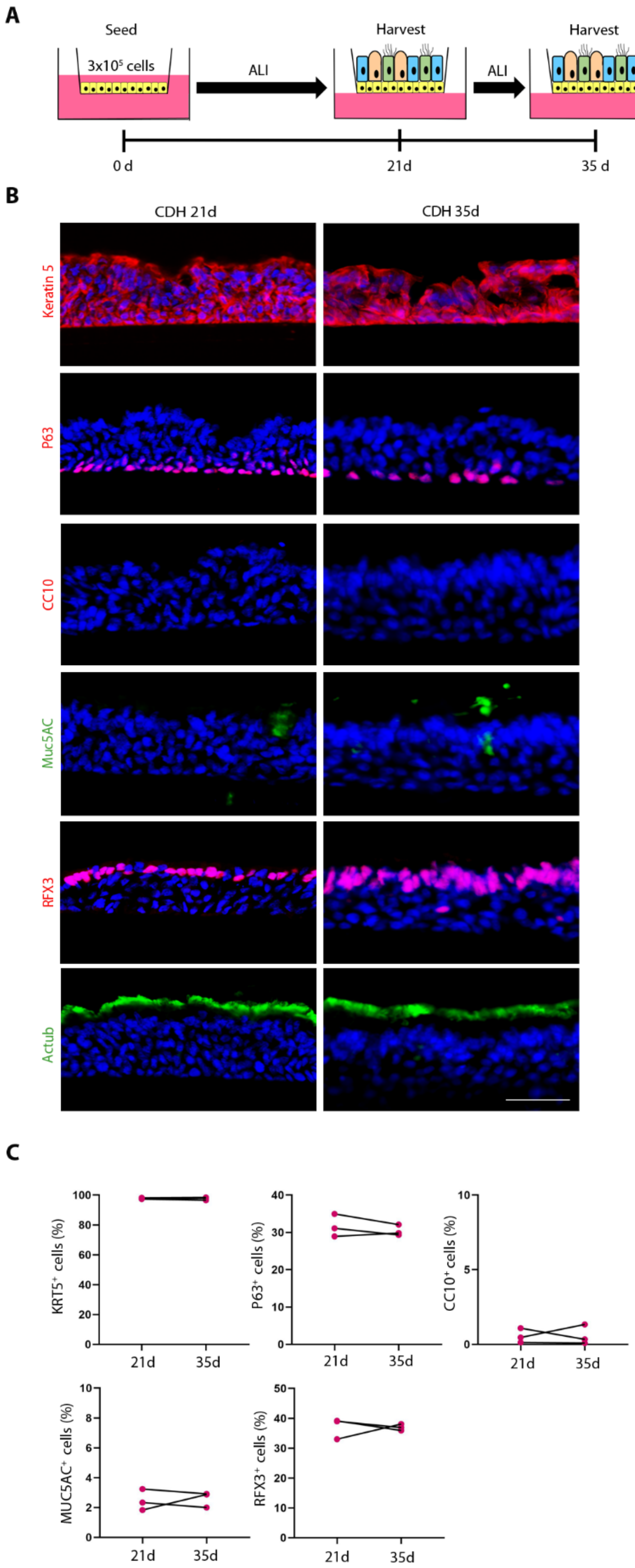


**Fig. S4. Prolonged culture in ALI has no effect on abnormal differentiation of the 7 CDH BSC lines.** (**A**) Schematic of the workflow. (**B**) Representative fluorescence images of antibody staining for epithelial markers in ALI cultures of CDH BSCs at day 21 and day 35 (n=3 lines). Nuclei were stained by DAPI. (**C**) The relative abundance of each labelled cell type was quantified for each condition in triplicates. Each dot represents one control BSC line. At least 3 technical replicates for each line were evaluated. Scale bar, 25 μm.

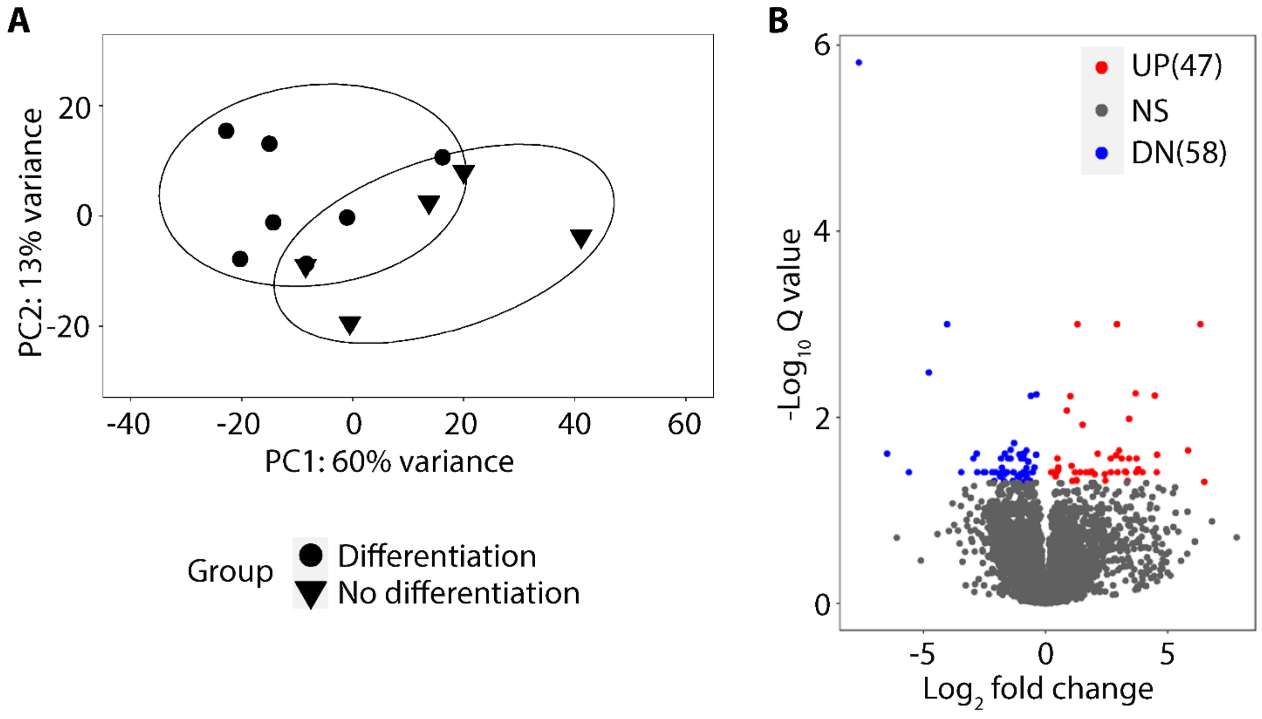


**Fig. S5. Transcriptome profile is not sufficient to separate the two subgroups of CDH BSCs that differ in epithelial differentiation capacity.** (**A**) Principal component analysis of bulk RNA-seq results of the 7 TA BSCs that were able to differentiate and the 5 CDH BSCs that were unable to differentiate in ALI. (**B**) Volcano plot that highlights differentially expressed genes comparing CDH BSCs that failed to differentiate to CDH BSCs that were able to differentiate. Genes with an adjusted p value <0.05 were marked. UP, upregulated. DN, downregulated.

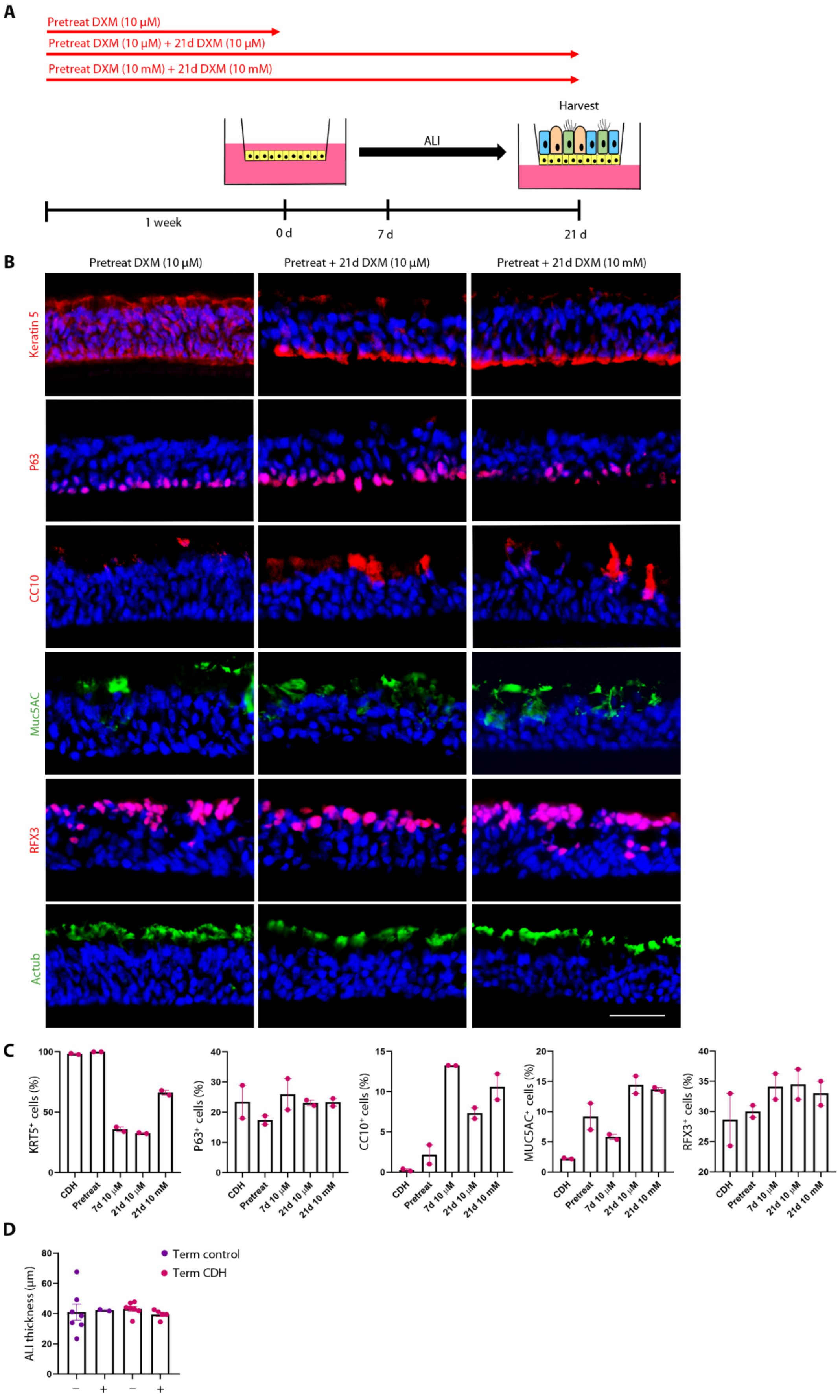


**Fig. S6. DXM treatment at two different concentrations and for various duration was tested for its effect on CDH BSC differentiation.** (**A**) Schematic of the workflow. DXM (10 µM and 10 mM) pretreatment alone during cell expansion and pretreatment + treatment in ALI were tested. (**B**) Representative fluorescence images of staining for epithelial markers under different treatment conditions.

Nuclei were stained with DAPI. Scale bar, 25 µm. (**C**) The relative abundance of each labelled cell type was quantified for each condition in triplicates. Each dot represents one CDH BSC line. Results of the untreated and pre-treatment + day 7 conditions, which were shown in Fig. 5F, were included for direct comparison to other tested conditions. (**D**) Quantification of the thickness of the epithelial layer in day 21 ALI culture of control and CDH BSCs with and without DXM treatment. Bar graph represents mean ± SEM for each experimental group. At least 3 technical replicates for each line were evaluated.

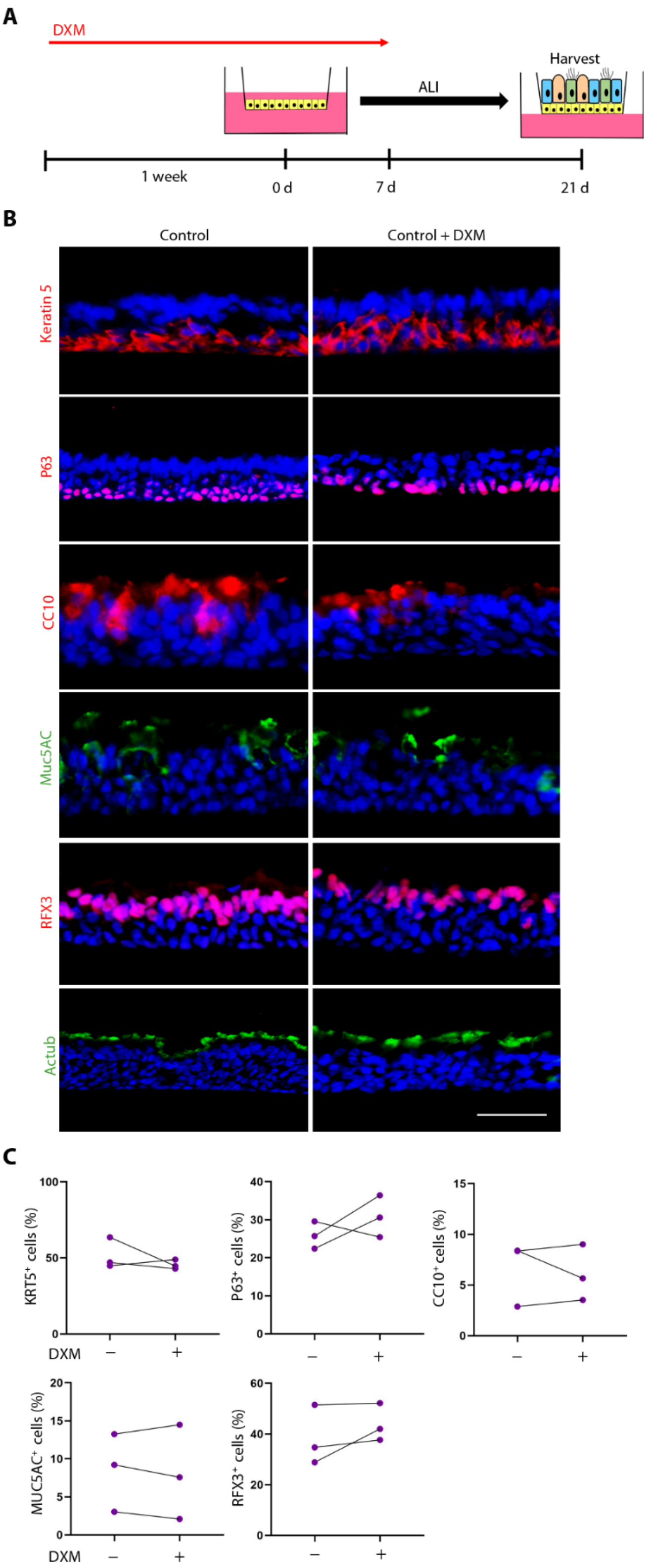


**Fig. S7. DXM treatment has no effect on epithelial differentiation of non-CDH control BSCs in ALI.** (**A**) Schematic of the workflow. (**B**) Representative fluorescence images of antibody staining for epithelial markers in untreated and treated ALI cultures of control BSCs (n=3 lines). Nuclei were stained by DAPI. Scale bar, 25 μm. (**C**) The relative abundance of each labelled cell type was quantified for each condition in triplicates. Each dot represents one control BSC line.

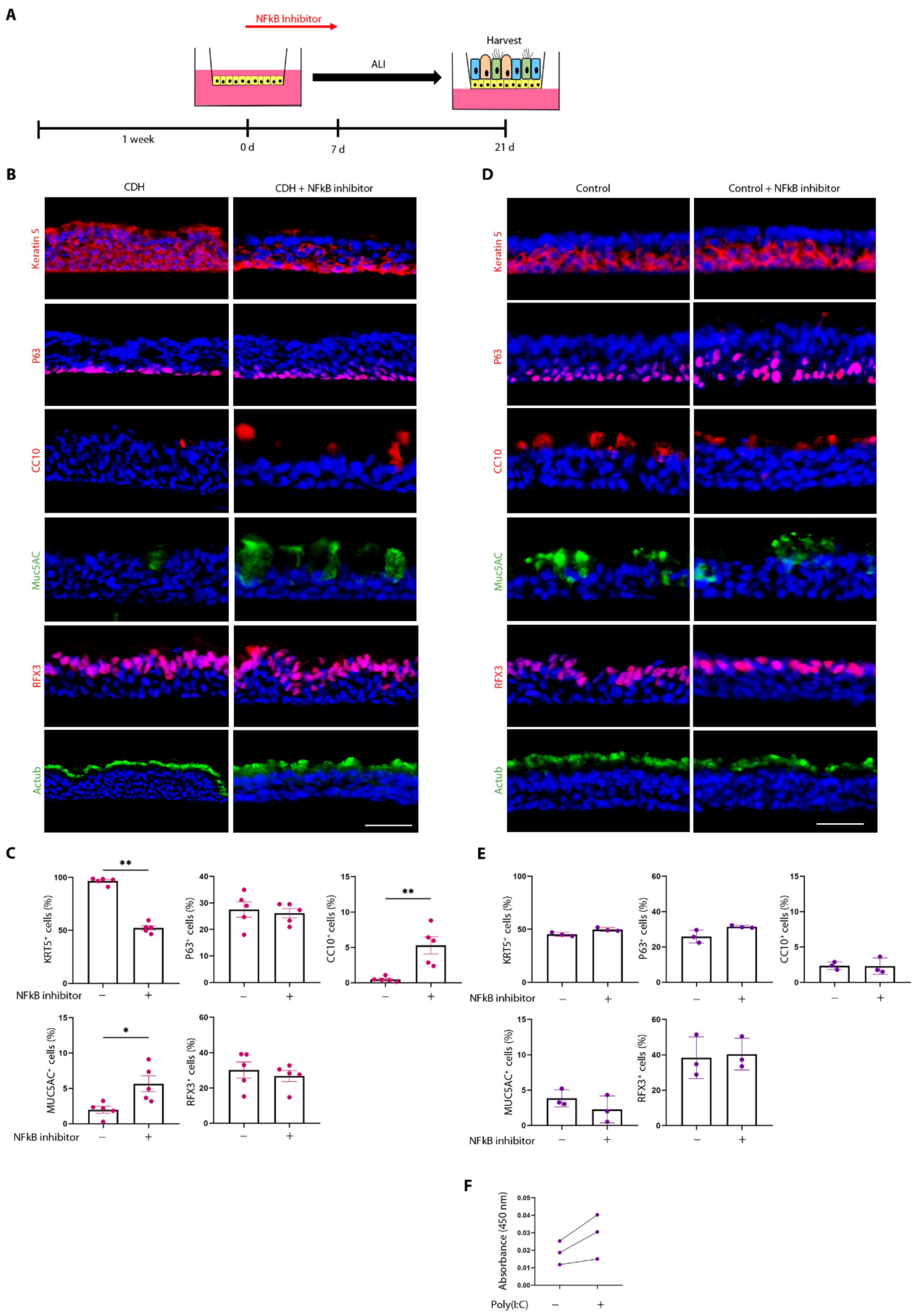


**Fig. S8. NF-κB inhibitor rescues epithelial differentiation defects of CDH BSCs and has no effect on differentiation of control BSCs in ALI.** (**A**) Schematic of the workflow. (**B**) Representative fluorescence images of antibody staining for epithelial markers in untreated and treated ALI cultures of CDH BSCs (n=5 lines). Nuclei were stained by DAPI. Scale bar, 25 μm. (**C**) The relative abundance of each labelled cell type was quantified for each condition in triplicates. Each dot represents one CDH BSC line. (**D**) Representative fluorescence images of antibody staining for epithelial markers in untreated and treated ALI cultures of control BSCs (n=3 lines). Nuclei were stained by DAPI, 25 μM. (**E**) The relative abundance of each labelled cell type was quantified for each condition in triplicates. Each dot represents one control BSC line. Bar graph represents mean ± SEM for each experimental group. At least 3 technical replicates for each line were evaluated. (**F**) Absorbance (450 nm) measurements of DNA binding activity of nuclear NF-κB by ELISA in untreated and Poly(I:C) treated control BSCs (n=3 lines). *p<0.05 and **p<0.01 by Mann–Whitney U test. Scale bar,

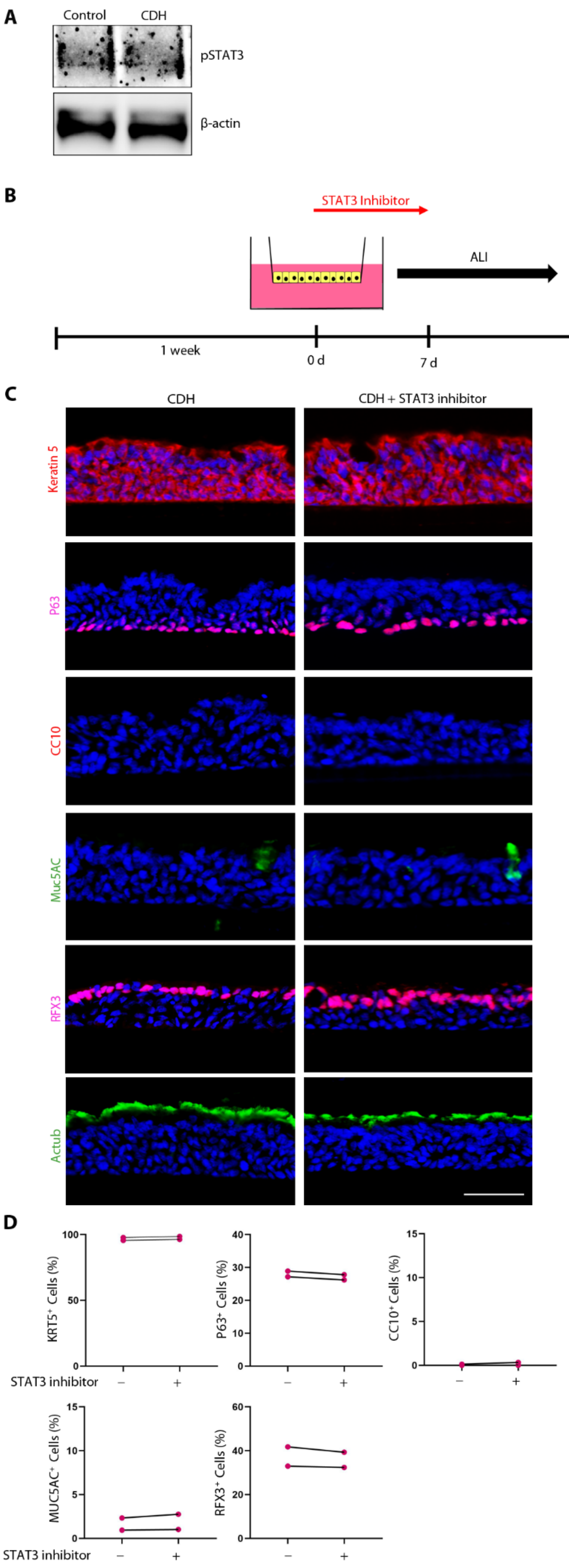


**Fig. S9. STAT3 inhibition had no effect on ALI differentiation phenotype of CDH BSCs.** (**A**) Representative Western Blot for phosphorylated STAT3 (pY705) comparing control and CDH BSCs. (**B**) Schematic of the workflow. STAT3 inhibitor S3I-201 (20 μM) was added for the first week during ALI differentiation of CDH BSCs (n=2 lines) (**C**) Representative fluorescence images of staining for epithelial markers. Nuclei were stained by DAPI. Scale bar, 25 µm. (**D**) The relative abundance of each labelled cell type was quantified in triplicates and shown for each line with and without treatment. More than 6,000 cells were counted using stained sections of ALI cultures for each line.

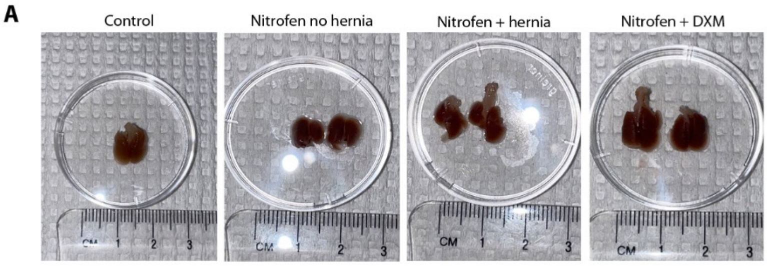


**Fig. S10. Prenatal dexamethasone treatment partially rescues the lung size in the nitrofen rat model of CDH.** (**A**) Macroscopic comparison of the lung size within each group.

### DETAILED MATERIALS AND METHODS

#### Study design

The study was designed to characterize TA-derived BSCs from CDH and non-CDH newborns and compare their gene expression and differentiation capacity *in vitro*. It is considered unethical to obtain lung biopsies from CDH patients. Therefore, we provide a novel method to use TA-derived BSCs as a surrogate to model proximal epithelial lung defects in CDH newborns. All experiments, patient enrollment, and sample acquisition were approved by the appropriate regulatory and animal welfare committees at Massachusetts General Hospital (IRB #2019P003296, PI: Lerou), Boston Children's Hospital (IRB #2000P000372, PI: High). After parental consent, TA samples were obtained at NICUs at Massachusetts General Hospital and Boston Children's Hospital during routine tracheal suctioning of intubated term CDH patients and term (37-40 weeks) and preterm (24-28 weeks) control patients (Table 1 + Suppl. Table 1). All samples were obtained similarly within 24-48 h after birth. TAs were transferred to the laboratory on ice and processed according to our standardized protocol *(66)* immediately after arrival. Human fetal lung sections were obtained from a previously established lung tissue bank at the University of Manitoba (IRB protocol #HS15293, PI: Keijzer) *(25)*. All assays were run in a minimum of triplicates and all data points including outliers are shown in the figures.

#### Nitrofen rat model of CDH

All animal work was approved by institutional animal care and use committee at Massachusetts General Hospital (IACUC #2022N000003, PI: Ai). Timed pregnant Sprague-Dawley rats were purchased from Charles River Laboratory and were randomly assigned to the respective treatment groups. Rat dams at E9.5 were anesthetized with isoflurane and orally gavaged with either nitrofen (100mg in 1 mL olive oil) or olive oil alone (vehicle) *(24)*. Rat fetuses were harvested at E21.5.

#### BSC isolation, expansion, and epithelial differentiation in air liquid interface (ALI) culture

Derivation of BSCs from TA samples, BSC expansion, and differentiation in ALI were previously described *(37, 66)*. Briefly, cells in TA were pelleted by centrifugation and then plated into precoated T25 flasks in 4 mL small airway epithelial growth medium (SAGM, PromoCell, Cat#C-21070). Medium was changed every two days for 2-3 weeks until cells reached confluency (approx. $2.3x10^6$ cells). The culture was trypsinized (trypsin-EDTA 0.05%, Thermo Scientific, Cat#25200114) and expanded in a T75 flask and a

T175 flask (~15x$10^6$ cells) as P2. For each passage from P2, culture was split at 1:6. BSCs were cultured on coverslips and fixed for 15 min with 4% paraformaldehyde in PBS before immunostaining according to a standard protocol using the following primary antibodies: rabbit anti-P63 (1:100, Cat#13109S, Cell Signaling Technology), rabbit anti-KRT5 (1:100, Cat#ab53121, Abcam), and rabbit anti-NKX2.1 (1:100, Cat#sc-13040, Santa Cruz Biotechnology).

To induce BSC differentiation in ALI, 3x$10^5$ cells were seeded onto 0.4 µm Transwell membranes (Cat#3460, Corning) that were precoated with 804G-conditioned medium. Differentiation in Pneumacult medium (StemCell Technology, Cat#05001) started with removal of medium at the top chamber of the insert. Medium was changed daily during the first week of ALI and every other day during the following 2 weeks. At day 21, ALI cultures were assessed with transepithelial electrical resistance (TEER > 500 Ohm) with an epithelial Voltohmmeter (World Precision Instruments, Inc., Sarasota, FL) and then fixed for 15 min with 4% paraformaldehyde in PBS before they were processed for cryo-sectioning at a thickness of 14 µm. Sections of ALI cultures were immunostained according to a standard protocol using the following primary antibodies: rabbit anti-KRT5 (1:100, Cat#ab53121, Abcam), rabbit anti-P63 (1:100, Cat#13109S, Cell Signaling Technology), rabbit anti-CC10 (1:100, Cat#HPA031828, Sigma-Aldrich), mouse anti-Muc5AC (1:100, Cat#MA5-12178, Thermo Scientific), rabbit anti-RFX3 (1:100, Cat#HPA035689, Sigma Aldrich) and mouse anti-Actub (1:100, Cat#T6793, Sigma Aldrich) *(34, 37)*. RFX3 and Actub both label ciliated cells. We used the nuclear marker RFX3 for quantification of ciliated cells, while Actub was stained to confirm the polarity of the differentiated epithelium. For treatment with DXM (Cat#D4902, Sigma Aldrich), BSCs were treated for 7 days during cell expansion and for 7-21 days during ALI differentiation at 10µM and 10mM. For treatment with Poly (I:C) (10µM, Cat#tlrl-pic, Invivogen), BSCs were treated every 2 days during 7-day cell expansion followed by treatment at days 3, 5, 7, and 14 in ALI. For treatment with a specific NF-κB inhibitor JSH-23 (10 µM, Cat#S7351, Selleck Chemicals) or STAT3 inhibitor S3I-201 (20µM, Cat#S1155, Selleck Chemicals), ALI cultures were treated daily during the first week of differentiation.

**Transcriptome analysis of BSCs by bulk RNA-seq**

Bulk RNA-seq was performed by ActiveMotif.inc (Carlsbad, CA, USA). For RNA sequencing, 1x$10^6$ BSCs were centrifuged and cryopreserved in 50% FBS/40% SAGM/10%DMSO solution before further processing. Cellular RNA extraction was performed using Qiagen RNA RNeasy Kit (QIAGEN, Cat#74104). RNA was subjected to library preparation with Illumina TruSeq V2 kit (Illumina, Cat#20020594). Libraries were sequenced on Illumina NextSeq 500 as paired-end 42-nt reads. Reads were aligned using STAR version 2.6.0a to the UCSC GRCh38 genome build and counts were calculated against the corresponding gene annotation using htseq-count version 0.10.0.17,18. R package DESeq2 version 1.30.0 was used for differential expression analysis. Only genes with an average count above 2 were kept for further analysis. Differential expression analysis and principal component analysis were performed using the DESeq2 R package (1.30.0). Genes with an adjusted p value less than 0.05 (padj<0.05) were considered as differentially expressed genes (DEGs). DEGs were ranked by stat to perform Gene set enrichment analysis (GSEA) in GSEA Molecular Signatures Database using R package fgsea version 1.16.0. Heatmap was made using R package ComplexHeatmap version 2.6.2. Dot plot, volcano plot, and waterfall plot were generated using R package ggplot2 version 3.3.3. The pre-existing bulk RNA-seq dataset (GSE164358) of neonatal basal cells from nasopharyngeal (NP) aspirates was re-analyzed and used as control to compare the basal stem cell markers' expression level with our BSCs from tracheal aspirates *(38)*.

**Chromatin accessibility analysis of BSCs by ATAC-seq**

BSCs ($1x10^5$) were cryopreserved in 50% FBS/40% SAGM/10%DMSO solution prior to ATAC-seq by ActiveMotif.inc (Carlsbad, CA, USA). Cells were then thawed in a 37℃ water bath, pelleted, and tagmented using Nextera Library Prep Kit (Illuminutesa, Cat#FC-130-1064) as previously described *(67, 68)*. Tagmented DNA was purified using the MinutesElute PCR purification kit (Qiagen, Cat#13323), amplified with 10 cycles of PCR, purified using Agencourt AMPure SPRI beads (Beckman Coulter), and sequenced on Illumina NextSeq 500 sequencer as paired-end 42-nt reads. The paired-end 42 bp sequencing reads were then mapped to the human hg38 reference genome using the BWA version 0.7.12 algorithm with default settings. Only reads that pass Illumina's purity filter, align with no more than 2 mismatches, and map uniquely to the genome are used in the subsequent analysis. In addition, duplicate reads are removed. Alignments were extended in silico at their 3'-ends to a length of 200 bp and assigned to 32-nt bins along the genome. The resulting histograms (genomic "signal maps") were stored in bigWig files. Peaks were identified using the MACS version 2.1.0 algorithm at a cutoff of p-value 1e-7, without control file, and with the –nomodel option. Peaks that were on the ENCODE blacklist of known false ChIP-

Seq peaks were removed. Paired reads landing in peaks are counted using R package Rsubread version 2.6.4. Differential changes of ATAC-seq signal within peaks were performed using R package DESeq2 version 1.30.0. Motif analysis within the differential regions was performed by findMotifsGenome.pl function in HOMER version 4.11. R package karyoploteR version 1.16.0 was used to create karyoplot showing ATAC-seq tracks.

**Immunohistochemistry of human and rat lung tissue**

Tracheas and lungs were fixed in 4% paraformaldehyde/PBS and processed for paraffin embedding and sectioning before immunohistochemistry using standard staining procedures.

Paraffin sections (5 μm) of lung tissue samples from term CDH and non-CDH human fetuses and randomly chosen E21.5 rat fetuses were prepared and immunostained according to standard protocols using the following primary antibodies: rabbit anti-KRT5 (1:100, Cat#ab53121, Abcam), rabbit anti-P63 (1:100, Cat#13109S, Cell Signaling Technology), rabbit anti-CC10 (1:100, Cat#HPA031828, Sigma-Aldrich), mouse anti-Muc5AC (1:100, Cat#MA5-12178, Thermo Scientific), and rabbit anti-RFX3 (1:100, Cat#HPA035689, Sigma Aldrich).

**Western Blot and ELISA**

$2x10^6$ BSCs treated with or without 10μM dexamethasone for 1 week in culture, were lysed in 250 μL RIPA buffer with protease and phosphatase inhibitor. 20 μL of protein lysate per sample were loaded and the assay was performed according to standard protocols with primary antibodies against phosphorylated p65 subunit at Ser536 (1:1000, Cat#3033T, Cell signaling Technology), phosphorylated STAT3 at Tyr705 (1:1000, Cat#9145S, Cell Signaling Technology), and β-actin (1:2000, Cat#A5441, Sigma Aldrich).

Nuclear contents of $2x10^6$ CDH BSCs and control BSCs were extracted using phosphatase and protease inhibitors according to standard protocols (Cat#10009277, Cayman Chemicals). 10 μL of nuclear lysate per sample were loaded and the ELISA assay was performed according to the manufacturer`s protocol (Cat#10007889, Cayman Chemicals). Absorbance was read at 450 nm.

**Imaging and quantification**

Stained slides were examined with a Nikon Ti inverted fluorescence microscope or Zeiss AX10 brightfield microscope. All images were processed using ImageJ software. Quantification was performed by two independent and blinded examiners as follows. For ALI and lung sections, 5 individual images per cell line at 20X were randomly selected and captured. DAPI-positive nuclei or hematoxylin-labelled nuclei in airway epithelium were counted as the total number of cells per 20X image. Cells labelled with specific epithelial markers in the same image were counted and reported as the percentage of the total cell population.
